# Enhanced stability of the SARS CoV-2 spike glycoprotein trimer following modification of an alanine cavity in the protein core

**DOI:** 10.1101/2022.11.08.515567

**Authors:** Pantelis Poumbourios, Christine Langer, Irene Boo, Tasnim Zakir, Rob J. Center, Anouschka Akerman, Vanessa Milogiannakis, Anupriya Aggarwal, Stuart Turville, Heidi E. Drummer.

## Abstract

The spike (S) glycoprotein of SARS CoV-2 is the target of neutralizing antibodies (NAbs) that are crucial for vaccine effectiveness. The S1 subunit binds ACE2 while the S2 subunit mediates virus-cell membrane fusion. S2 is a class I fusion glycoprotein and contains a central coiled coil that acts as a scaffold for the conformational changes associated with fusion function. The coiled coil of S2 is unusual in that the 3-4 repeat of inward-facing positions are mostly occupied by polar residues that mediate few inter-helical contacts in the prefusion trimer. We examined how insertion of bulkier hydrophobic residues (Val, Leu, Ile, Phe) to fill a cavity formed by Ala^1016^ and Ala^1020^ that form part of the 3-4 repeat affects the stability and antigenicity of S trimers. Substitution of Ala^1016^ with bulkier hydrophobic residues in the context of a prefusion-stabilized S trimer, S2P-FHA, was associated with increased thermal stability. The trimer stabilizing effects of filling the Ala^1016^/Ala^1020^ cavity was linked to improved S glycoprotein membrane fusion function. When assessed as immunogens, two thermostable S2P-FHA mutants derived from the ancestral isolate, A1016L (16L) and A1016V/A1020I (VI) elicited very high titers of neutralizing antibodies to ancestral and Delta-derived viruses (1/2,700-1/5,110), while neutralization titer was somewhat reduced with Omicron BA.1 (1/210-1,1744). The antigens elicited antibody specificities that could compete with ACE2-Fc for binding to the receptor-binding motif (RBM) and NAbs directed to key neutralization epitopes within the receptor-binding domain (RBD), N-terminal domain (NTD) and stem region of S2. The VI mutation enabled the production of intrinsically stable Omicron BA.1 and Omicron BA.4/5 S ectodomain trimers in the absence of an external trimerization motif (T4 foldon). The VI mutation represents a method for producing an intrinsically stable trimeric S ectodomain glycoprotein vaccine in the absence of a foreign trimerization tag.

**AUTHOR SUMMARY:** First-generation SARS CoV-2 vaccines that generate immune responses to ancestral Spike glycoprotein sequences have averted at least 14.4 million deaths, but their effectiveness against the recently emerged Omicron lineages is reduced. The updating of booster vaccines with variant Spike sequences are therefore likely required to maintain immunity as the pandemic continues to evolve. The Spike is a trimeric integral membrane protein with a membrane spanning sequence at its C-terminus. The Spike protein-based vaccine that is currently licensed for human use is produced by a complex process that reconstitutes the Spike in an artificial membrane. Alternatively, production of the Spike trimer as a soluble protein generally requires replacement of the membrane spanning sequence with a foreign often highly immunogenic trimerization motif that can complicate clinical advancement. We used systematic structure-directed mutagenesis coupled with functional studies to identify an alternative stabilization approach that negates the requirement for an external trimerization motif or membrane-spanning sequence. The replacement of 2 alanine residues that form a cavity in the core of the Spike trimer with bulkier hydrophobic residues resulted in increased Spike thermal stability. Thermostable Spike mutants retained major conserved neutralizing antibody epitopes and the ability to elicit broad and potent neutralizing antibody responses. One such mutation, referred to as VI, enabled the production of intrinsically stable Omicron variant Spike ectodomain trimers in the absence of an external trimerization motif. The VI mutation potentially enables a simplified method for producing a stable trimeric S ectodomain glycoprotein vaccine.

---

The SARS CoV-2 betacoronavirus has led to the death of more than 6.4 million people with a strong age dependent fatality rate. First-generation vaccines that deliver ancestral SARS CoV-2-derived viral Spike glycoprotein (S) sequences for its in vivo expression and neutralizing antibody (NAb) induction have been rolled out across the globe and have proven highly effective at preventing symptomatic and severe COVID-19. The viral S glycoprotein mediates receptor attachment and virus-cell membrane fusion and is the target of NAbs, which play a critical role in protection against SARS CoV-2 transmission and disease progression (1–4). The mature spike comprises 2 functional subunits, S1 and S2, that are derived from a polyprotein precursor, S, by furin cleavage of an oligobasic motif as it transits the Golgi. ACE2 receptor attachment is mediated by the RBD within the large subunit, S1, while membrane fusion is mediated by the small subunit, S2, which contains the fusion peptide. S1 and S2 form a heterodimer via non-covalent interactions. S1-S2 is assembled into a higher-order trimeric structure with a coiled coil-forming α-helix of S2 (amino acids 986-1033; referred to as CH) forming the core of the trimer (5) **(Fig. 1A)**. A membrane-spanning sequence at the C-terminus of S2 stabilizes the trimer and anchors it to the viral or cell membrane (6). The ACE2 RBD sits atop the S1 glycoprotein trimer and presents in “up” ACE2-binding-ready and “down” inert orientations (7). Following receptor attachment, S2 is cleaved by the TMPRSS2 protease at the cell surface or by cathepsin L following endocytosis to liberate the fusion peptide of S2 and enable full fusion activation (For review: (8). The S glycoprotein mediates membrane fusion via a class I mechanism whereby an activation trigger (ACE2-binding by S1, TMPRSS2 cleavage of S2) causes the S2 subunit of the metastable pre-fusion trimer to refold into a stable trimer of hairpins, bringing the N-terminal fusion peptide and C-terminal membrane spanning sequences together such that their associated membranes fuse (9).

**Fig. 1.**
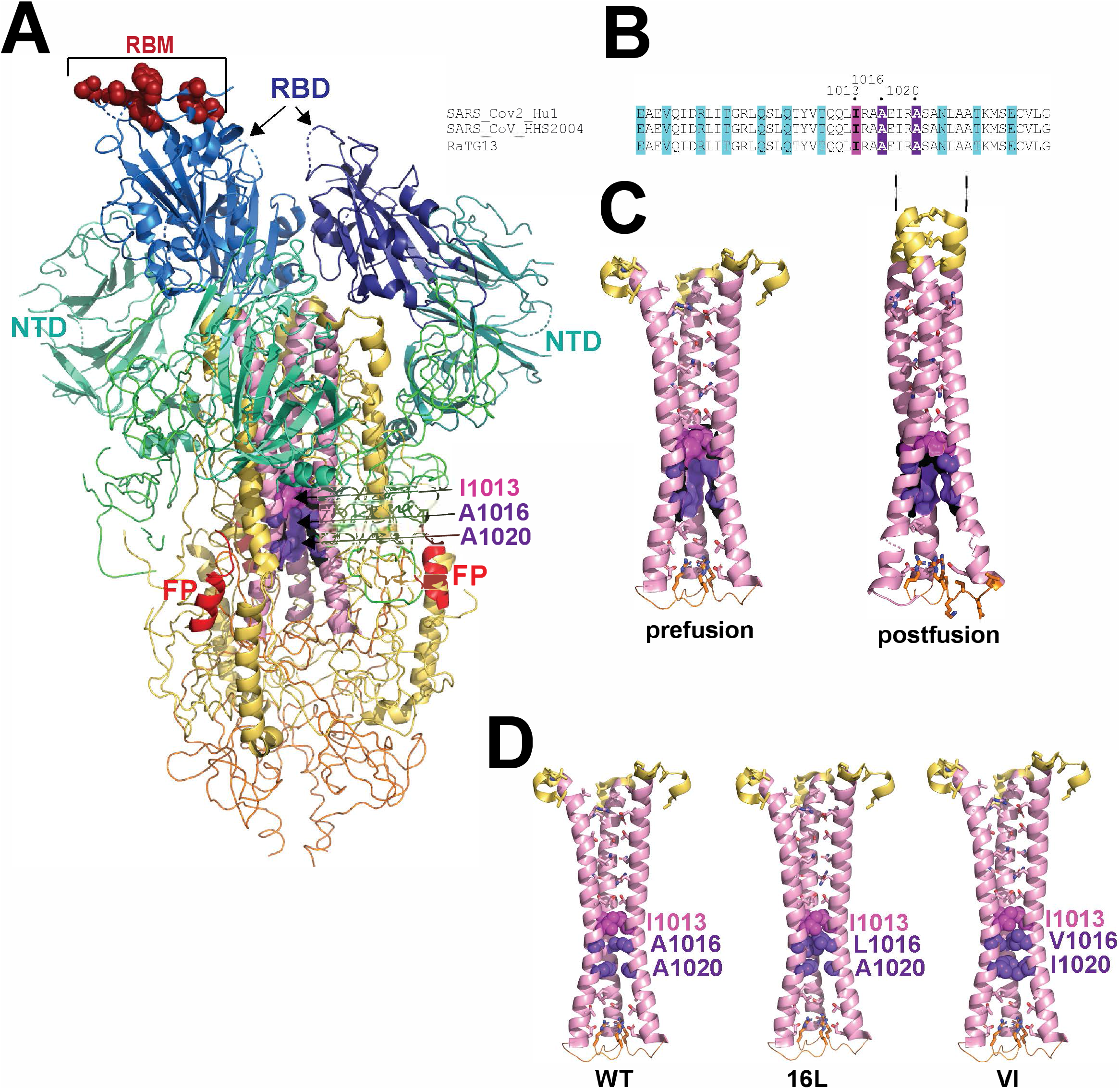
**A**, Three-dimensional structure of the SARS CoV-2 S ectodomain drawn with the coordinates PDB ID 6VSB (4). S1 is shown in green and blue, RBM, ACE2 receptor binding motif with receptor-interacting amino acids in red; RBD, receptor binding domain in blue; NTD, N-terminal domain in teal. S2 is shown in yellow and pink: FP, fusion peptide in red; central coiled coil in pink with Ala cavity highlighted in purple. **B**, a comparison of the central coiled coil in the prefusion (PDB ID 6VSB (4)) and postfusion (PDB 6XRA (9)) conformations. Although incompletely shown here, HR1 helices (C-terminal part shown in yellow) extend the coiled coil upward toward the cellular membrane in the postfusion conformation (9). **C**, heptad repeat motifs within the coiled coil sequences of 3 betacoronaviruses. **D**, Illustration of how the Ala cavity of prefusion S might be filled following substitution with hydrophobic amino acids. 16L and VI are codes for A1016L and A1016V/A1020I mutations.

The sites of vulnerability to NAbs within S have been revealed with the isolation of monoclonal NAbs (mNAbs) from COVID-19 patients and vaccinees. The RBD is an immunodominant antibody target in natural infection (10) and highly potent NAbs directed to the RBD can block infection by binding to the ACE2 receptor-binding motif and directly blocking ACE2 binding, via steric blockade of ACE2 binding, by locking RBDs in the down orientation to preclude ACE2 binding, or by triggering premature refolding of S1-S2 into the post-fusion state with shedding of S1 (11–18). The NTD of S1 has also been identified as a supersite of vulnerability and comprises multiple antigenic sites (12, 15, 19–21). This region exhibits a high degree of plasticity, acquiring point mutations, deletions, insertions and glycan additions enabling antibody evasion (22). Conserved neutralization epitopes external to the RBD and NTD have also been identified including within the stem helix of S2 (12, 23, 24). Immunity induced by natural infection or vaccination is believed to be driving the emergence of variants of concern (VOCs) which primarily contain mutations in the RBD and NTD (25–28). Variants of concern are associated with successive waves of infections across the globe with Alpha, Delta and now Omicron infections, respectively, dominating the pandemic. Key mutations observed in the RBD of VOCs include K417T/N, N439K, N440K, L452R, T478K, E484K/Q/A, F486V and N501Y, while in the NTD, deletion of amino acids 24-26, 69-70, 142-144, 156-157 and 242-245 have been observed. COVID-19 vaccines based on ancestral SARS Cov-2 sequences proved to be highly effective against Alpha and Delta variants, however this was reduced with the emergence of the Omicron lineage of VOCs, which contains at least 30 mutations in S, including at least 15 mutations within the RBD (29–31). Reduced vaccine efficacy against Omicron lineage subvariants appears to correlate with reduced in vitro neutralization potency of vaccinee sera against these VOCs, especially when antibody levels wane over time (13, 32–35). Boosting with VOC-matched vaccines may help to maintain immunity against emerging viral variants as the COVID-19 pandemic evolves (34, 36, 37).

Class I viral fusion glycoproteins such as Env of retroviruses, HA of orthomyxoviruses and S of coronaviruses contain a central coiled coil, which acts as a scaffold for the conformational changes associated with the membrane fusion process (4, 5, 9, 38–45) (**Fig. 1A**). In the former 2 viral families, the inward-facing positions of the coiled coil are occupied by hydrophobic residues in a 3-4 repeat that stabilize the coiled coil via knob into-holes packing interactions. In the case of SARS CoV-2, these positions within the central coiled coil of S2 (formed by CH helices) are mostly occupied by polar residues that mediate few inter-helical contacts in the prefusion trimer (**Fig. 1B**). In the postfusion S trimer, the N-terminal 2/3 of the coiled coil is brought together by the packing of HR1 helices that extends the coiled coil in an N-terminal direction by 110Å. In this conformation the inward facing residues are close enough for hydrogen bonds to form **(Fig. 1C)**. Within the prefusion S coiled coil, Ile^1013^ is a point of contact between the 3 CH helices and forms a small hydrophobic core through inter-helical contacts with Ile^1013^ and with Leu^1012^. These interactions form a hydrophobic ceiling above a cavity formed by Ala^1016^ and Ala^1020^ that occupy central positions of the coiled coil (**Fig. 1A-D**). Classical studies on protein folding and stability revealed that cavities in a protein’s core are destabilising and filling the cavity with bulkier hydrophobic residues can improve thermal stability and function (46). In this study, we examined how filling the Ala^1016^ and Ala^1020^ cavity with bulkier hydrophobic amino acids influences protein expression, stability, antigenicity and immunogenicity in guinea pigs. Our study identified thermostable trimeric S mutants that elicit NAbs in small animals that maintain potency against Delta and Omicron VOCs. The cavity filling A1016V/A1020I mutation stabilized a trimer comprising the Omicron BA.1 and BA.4/5 S ectodomains and negated the requirement for an external motif to maintain trimerization. The A1016V/A1020I mutation represents a method for producing an intrinsically stable trimeric S ectodomain glycoprotein vaccine in the absence of a foreign trimerization tag.

## RESULTS

### Characterization of S2P-FHA

A CMV promoter driven expression vector was used to produce a soluble form of the S glycoprotein known as S2P (5), which comprises residues 16-1208 of the ancestral Hu-1 S glycoprotein, a furin cleavage site mutation, R^682^RAR-> G^682^SAS, and a di-Pro substitution at positions 986 and 987. T4 foldon, octa-His and avitag sequences were added to the C-terminus to give S2P-FHA. Following partial purification by divalent cation affinity chromatography, size exclusion chromatography (SEC) of the S2P-FHA protein revealed a major peak coeluting with thyroglobulin (669 kDa) that was collected as a homogenous protein as indicated by SEC and SDS-PAGE (**S1 Fig A, B**). A biotinylated form of purified S2P-FHA exhibited binding activity with hACE2-Fc, (human ACE2 residues 19-615 linked to the Fc domain of human IgG1) and various human mNAbs in ELISA (**S1 Fig C**). A thermofluor assay indicated that the S2P protein possessed relatively low thermal stability, exhibiting a melting temperature of 43.6°C (**S1 Fig D**).

### Effects of Ala cavity substitutions on thermostability

We asked how replacement of Ala^1016^ and Ala^1020^ with bulkier hydrophobic residues (Val, Leu, Ile, Phe) affects the stability of the SARS CoV-2 S trimer. (2 examples are shown in **Fig. 1D**). Alanine^1016^ and Ala^1020^ were singly and doubly substituted with Val, Ile Leu and Phe in S2P-FHA and the 293F-expressed glycoproteins partially purified by divalent cation affinity chromatography. SDS-PAGE indicated a range of yields: S2P, A1016V, A1020V, A1020I > A1016V/A1020V, A1016I, A1016L, A1020L, A1016V/A1020I > A1016I/A1020I, A1016L/A1020L, A1016V/A1020L, A1016V/A1020F, A1016I/A1020F > A1016F/A1020F (**Fig. 2A**). A thermofluor assay was used to examine the thermostability of S2P-FHA and the mutants. S2P-FHA comprised a major species with a melting temperature of 43.6°C and a more stable minor species with a melting temperature of 58°C (**Fig. 2B**). The substitution of hydrophobic residues into position 1016 increased the proportion of the 58°C species with A1016L being the most stabilizing mutation. By contrast, the 43.6°C species remained the predominant form with substitutions of A^1020^. Double substitutions were all associated with the stable form, except for A1016V/A1020V.

**Fig. 2.**
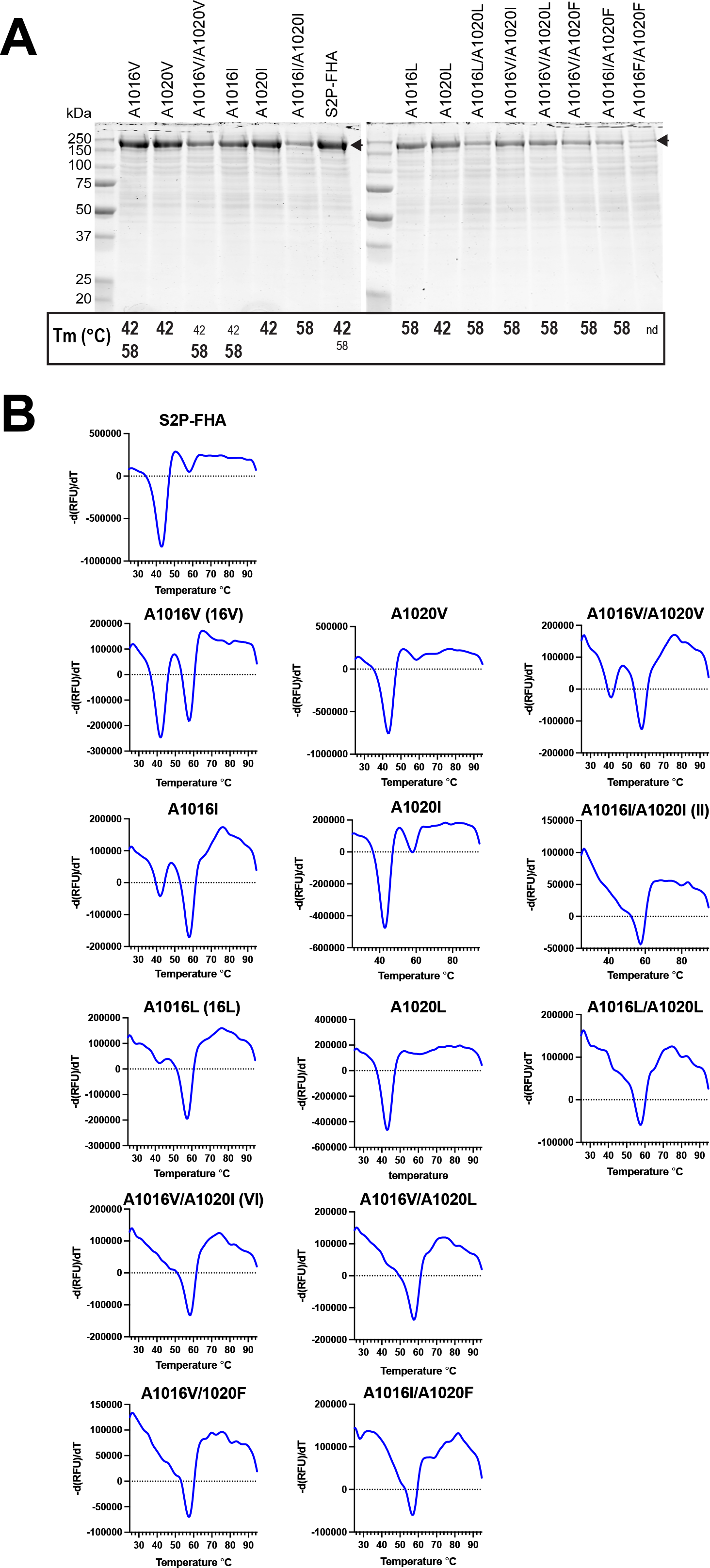
Expression and thermostability screen of Ala cavity mutants. **A**, Coomassie blue stained S2P-FHA mutants purified from culture supernatants with TALON resin following SDS-PAGE under reducing conditions. The melting temperatures (Tm) of S2P-FHA mutants obtained in **B** are indicated below each lane. Bold type: major species; small font: minor species; nd: not determined. **B**, Differential scanning fluorimetry of S2P-FHA proteins shown in A using SYPRO Orange. The rate of change of fluorescence over time [–d(RFU)/dt] as a function of temperature is shown. Graphs are representative of at least two independent experiments.

A subset of mutants with favourable thermostability/yield characteristics were purified to homogeneity and re-analysed in the thermofluor assay (**Fig. 3A-C**). The thermofluor data obtained with partially pure proteins were largely recapitulated with the purified trimers. Increasing proportions of the 58°C form relative to the 43.6°C form was observed for the mutants as follows: A1016V/A1020I (VI)=A1016I/A1020I (II) > A1016L (16L) > A1016V (16V) > S2P-FHA (**Fig. 3D**). Interestingly, this hierarchy of stability was inversely correlated to trimer yield (**Fig. 3C**).

**Fig. 3.**
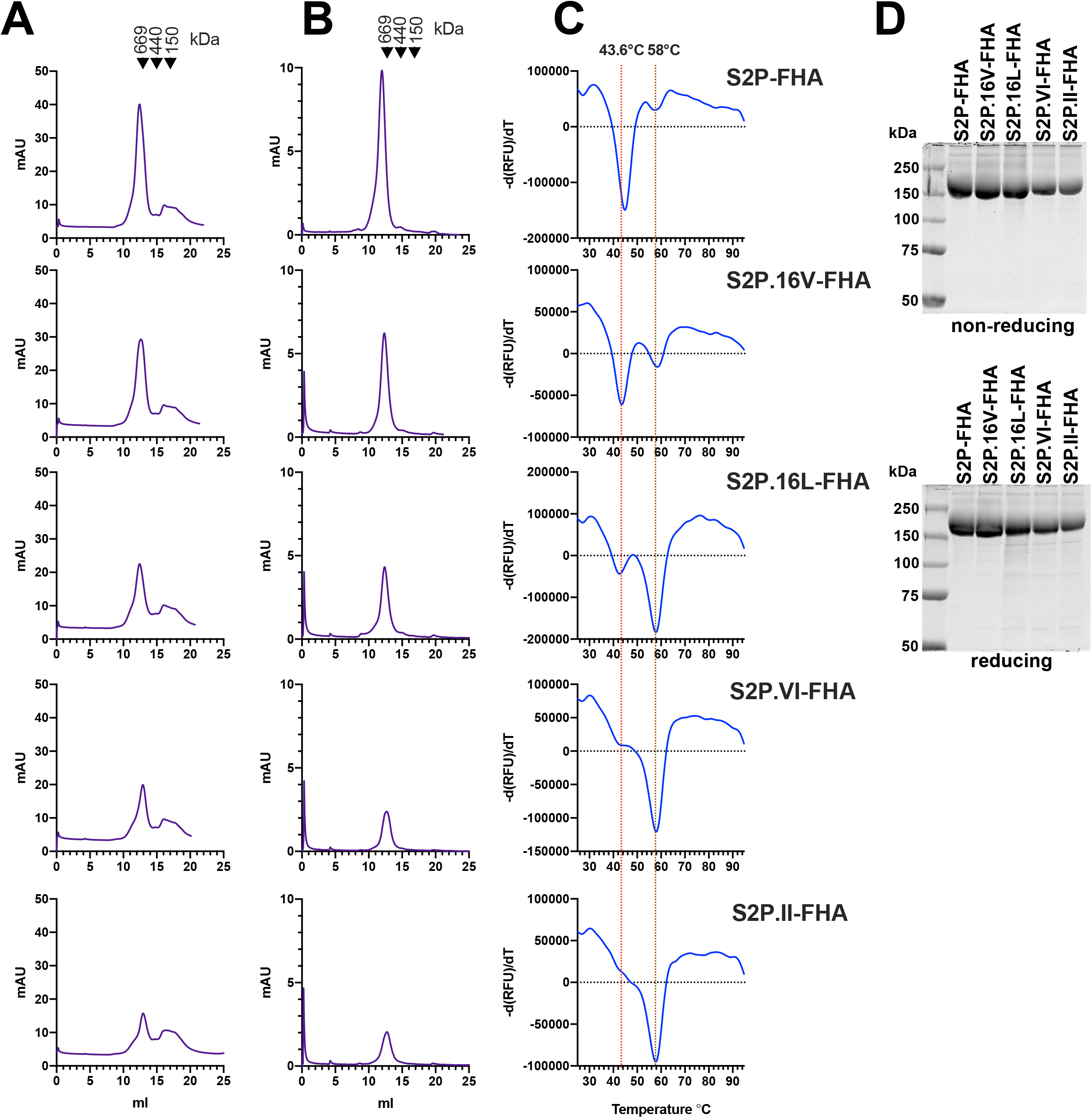
Purification and characterization of selected Ala cavity mutants. **A**, Superose 6 SEC of S2P-FHA proteins following elution from TALON resin. The calibration standards are thyroglobulin (669 kDa), ferritin, (440 kDa) and IgG (150 kDa). **B**, Superose 6 SEC of purified trimers following a freeze (−80°C)-thaw cycle. **C**, Differential scanning fluorimetry of purified S2P-FHA trimers. **D**, SDS-PAGE and Coomassie blue staining of purified proteins under nonreducing (top) and reducing (bottom) conditions.

### Functional properties of Ala cavity mutants

We next examined the effects of mutating Ala1016 and Ala1020 on the membrane fusion function of the ancestral SARS CoV-2 S glycoprotein. A1016V, A1016L, A1020V, A1020L and the VI mutations were introduced to the WH-Human1_EPI_402119 expression plasmid bearing codon-optimized full-length S. A western blot confirmed that the WT and mutated S glycoproteins were expressed and cleaved to S1 following transfection of 293T cells with the WH-Human1_EPI_402119 plasmids **(Fig. 4A)**. A cell-cell fusion assay was established to measure the membrane fusion function of the S glycoproteins. 293T effector cells were cotransfected with S expression vector, a bacteriophage T7 RNA polymerase expression plasmid (pCAG-T7) (47), and a furin expression plasmid (pcDNA3.1-Furin) (48). 293T-ACE2 target cells (49) were transfected with a T7 promoter-driven luciferase reporter plasmid (pTM*luc*) (50) and a TMPRSS2 expression plasmid (51). At 48 h post transfection, effector and target cells were cocultured for 3 h. A luciferase assay revealed that all S mutants possessed fusion activity that was between 1.5-2-fold higher than that of the WT **(Fig. 4B)**. Two control plasmids, S2P-1273, which expresses the full-length S glycoprotein containing Pro at positions 986 and 987 and a mutated cleavage site, and an HIV-1 Env expression plasmid, pcDNA3.1_AD8_-WT, lacked fusion activity. Consistent with the production of luciferase due to S-mediated fusion between effector and target cells, large multinucleated syncytia were observed for all S constructs but not for the controls S2P-1273 and HIV-1 Env **(Fig. 4C)**. The data indicate that Ala cavity filling mutations do not adversely affect membrane fusion function.

**Fig. 4.**
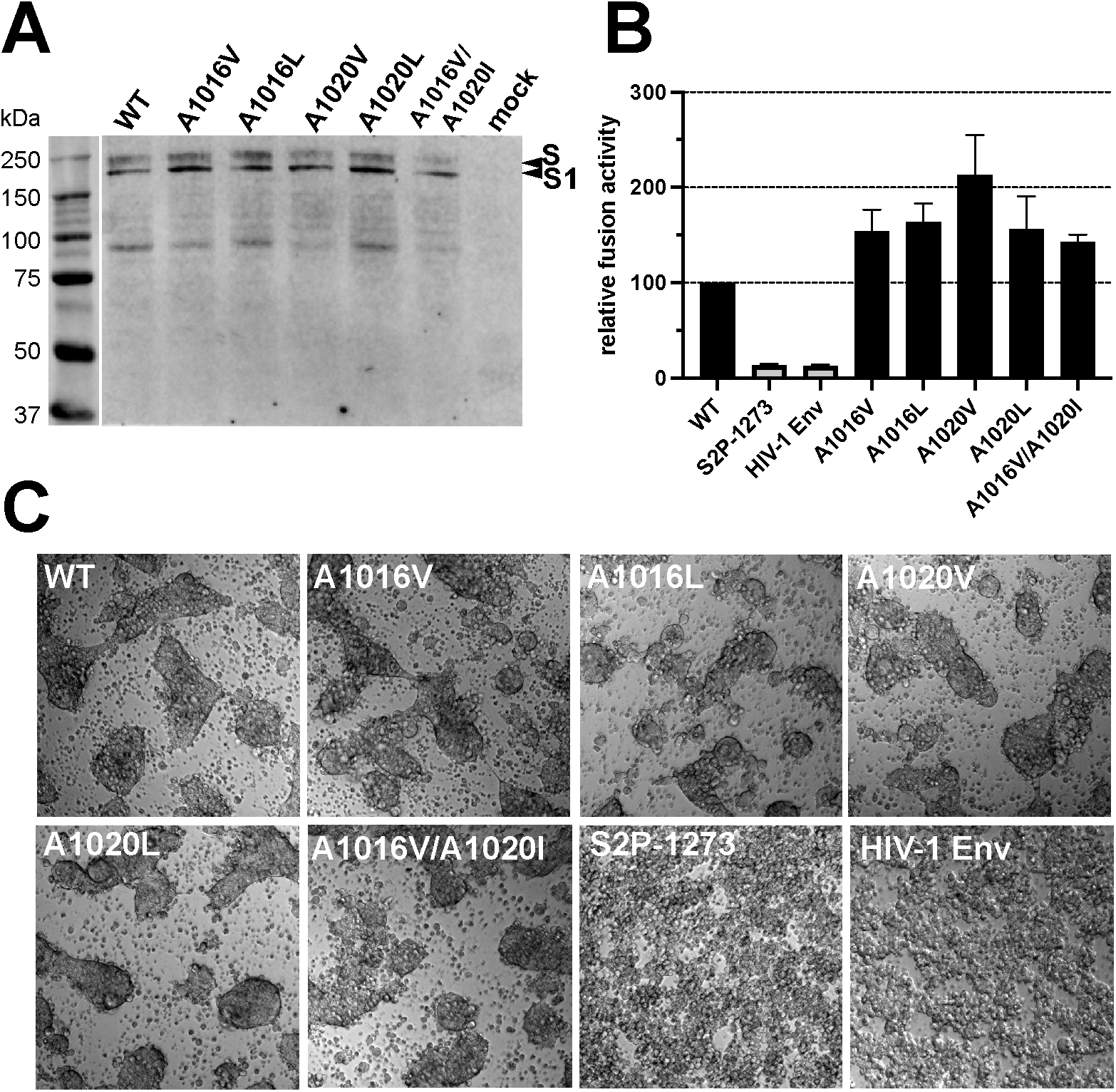
Effects of Ala cavity mutations on fusion function. **A**, Reducing SDS-PAGE and western blotting of SARS CoV-2 S glycoproteins expressed in 293T cells with rabbit anti-S1 polyclonal antibody. **B**, Cell-cell fusion activity of S glycoproteins determined in a luciferase reporter assay. Relative fusion activity: relative light units obtained with mutant and control vectors / relative light units obtained with WT S x 100. Mean ± SEM from 3 independent experiments shown. **C**, Representative microscopy fields at 10x magnification. Control vectors: S2P-1273 contains proline at positions 986 and 987 and lacks a furin cleavage site; HIV-1 Env: a HIV-1 glycoprotein expression vector, pcDNA3.1_AD8_-WT.

### Immunogenicity of Ala cavity mutants

Guinea pigs were used to examine whether the Ala cavity mutations can affect the magnitude and specificity of antibody responses to S2P-FHA trimers. Outbred guinea pigs were immunized with 30 μg of S2P-FHA, S2P.16L-FHA and S2P.VI-FHA in Addavax adjuvant at weeks 0, 4 and 14 and bleeds performed at weeks 6 and 16 (**Fig. 5A**). The neutralizing activity in vaccinal sera was determined using S-pseudotyped HIV luciferase reporter viruses and 293-ACE2 target cells as described previously (52). A comparison of week-6 and week-16 sera (bleeds taken 2 weeks after the 1^st^ and 2^nd^ boosts, respectively) using pseudotypes containing ancestral (Hu-1) S glycoprotein indicated potent neutralizing activity in S2P-FHA-, S2P.16L-FHA- and S2P.VI-FHA-immune sera with mean ID_50_s ranging from 1,700-1,900 for week-6 sera and 6,000-9,100 for week-16 sera. These data equate to ~3-5.4-fold increases in mean neutralization ID_50_ following the 2^nd^ boost, although statistical significance was not reached for S2P-FHA- and S2P.VI-FHA-immune sera (**Fig. 5B**). A capture ELISA, employing plate-bound avidin to capture biotinylated ancestral, Delta and Omicron BA.1 RBDs, was used to determine the RBD binding titers of vaccinal sera. **Figure 5C** shows no significant differences in binding profile between immunogen groups with geometric mean binding titers in the range 3.5×10^4^–1.36×10^5^. A small (~3-fold) but significant reduction in binding titer was observed for the Omicron BA.1 RBD for all 3 immunogen groups. Serum binding titers to ancestral, Delta and Omicron BA.1 S2P-FHA spike protein trimers bound directly to ELISA plates were next determined. **Figure 5D** indicates no significant differences in binding titers to the 3 S2P-FHA trimer variants for the 3 immunogen groups, which were in the range 3.0×10^5^-1.1×10^6^. These results suggest that the 3 immunogens generated similar titers of antibody able to bind the S2P-FHA spike trimer.

**Fig. 5.**
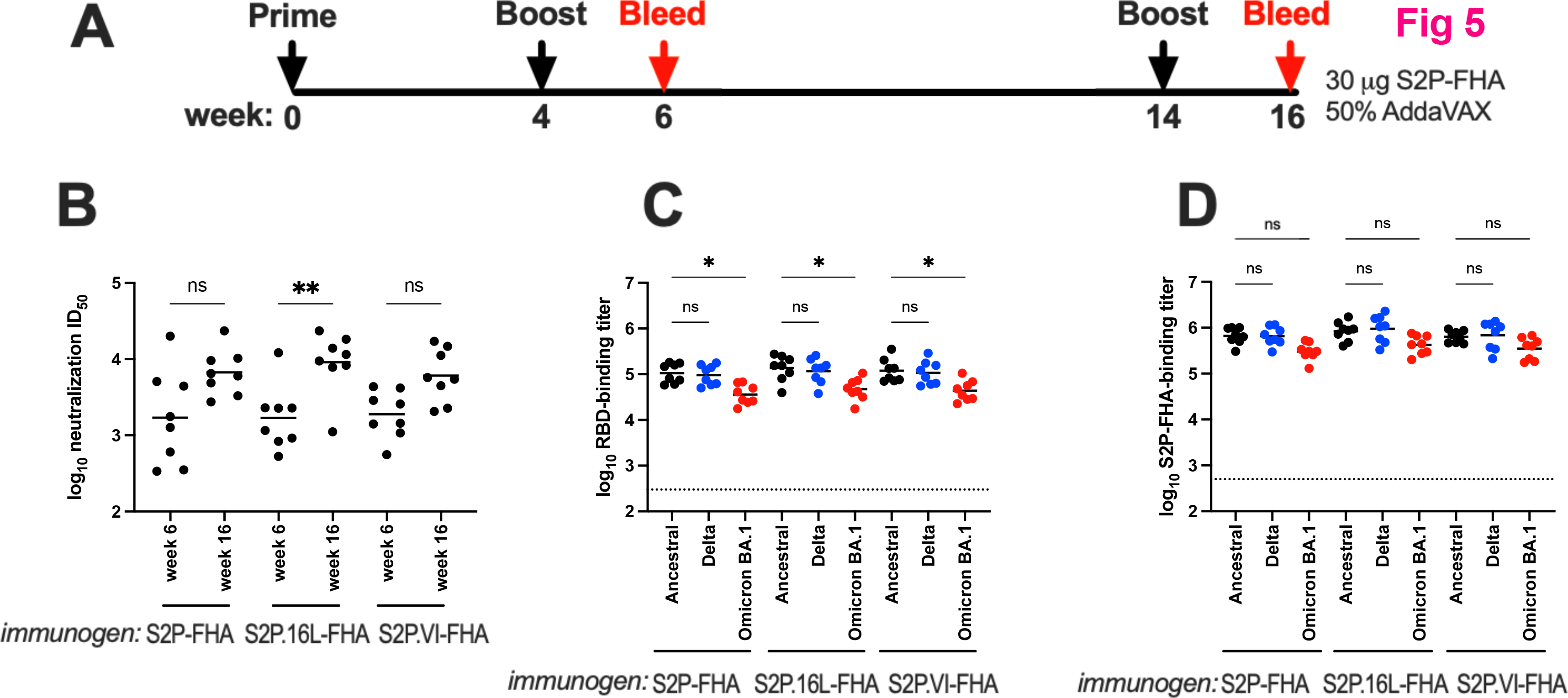
Immunogenicity of Ala cavity mutants. **A**, Immunization protocol. **B**, Pseudovirus neutralization ID_50_s of vaccinal sera obtained at weeks 6 and 16. The geometric mean pseudovirus neutralization ID_50_ of the control group of 8 animals receiving 3 doses of 50% Addavax-PBS was <1/120 at both time-points. **C and D**, ELISA binding titers of week-16 vaccinal sera to streptavidin-captured biotinylated RBDs or S2P-FHA trimers, respectively, derived from ancestral Hu-1, Delta and Omicron BA.1 (indicated below the graphs). The endpoint was determined as 5-times background optical density obtained in the absence of primary antibody. The horizontal bars are the geometric means. The horizontal dotted line is the geometric mean binding titer of the control group receiving 3 doses of 50% Addavax-PBS. A Kruskal-Wallis test was used to determine whether the differences in ID_50_s or binding titers are significantly different. ns, not significant; *, *P* < 0.05; **, *P* < 0.01.

We next assessed neutralizing activity against pseudotypes containing ancestral, Delta or Omicron BA.1 S glycoprotein and 293-ACE2 cells (52). Potent neutralizing activity against ancestral and Delta pseudoviruses was observed with S2P-FHA-, S2P.16L-FHA- and S2P.VI-FHA-immune sera with mean ID_50_s ranging from 3,900-5,100 (**Fig. 6A, S1 Table**). Serum neutralizing activities against pseudoviruses containing the Omicron BA.1 Spike were not significantly different relative to Hu-1 and Delta pseudovirus neutralization, although the IC_50_s against Omicron BA.1 pseudoviruses trended downwards with < 3-fold reductions in mean titer observed for the 3 groups.

**Fig. 6.**
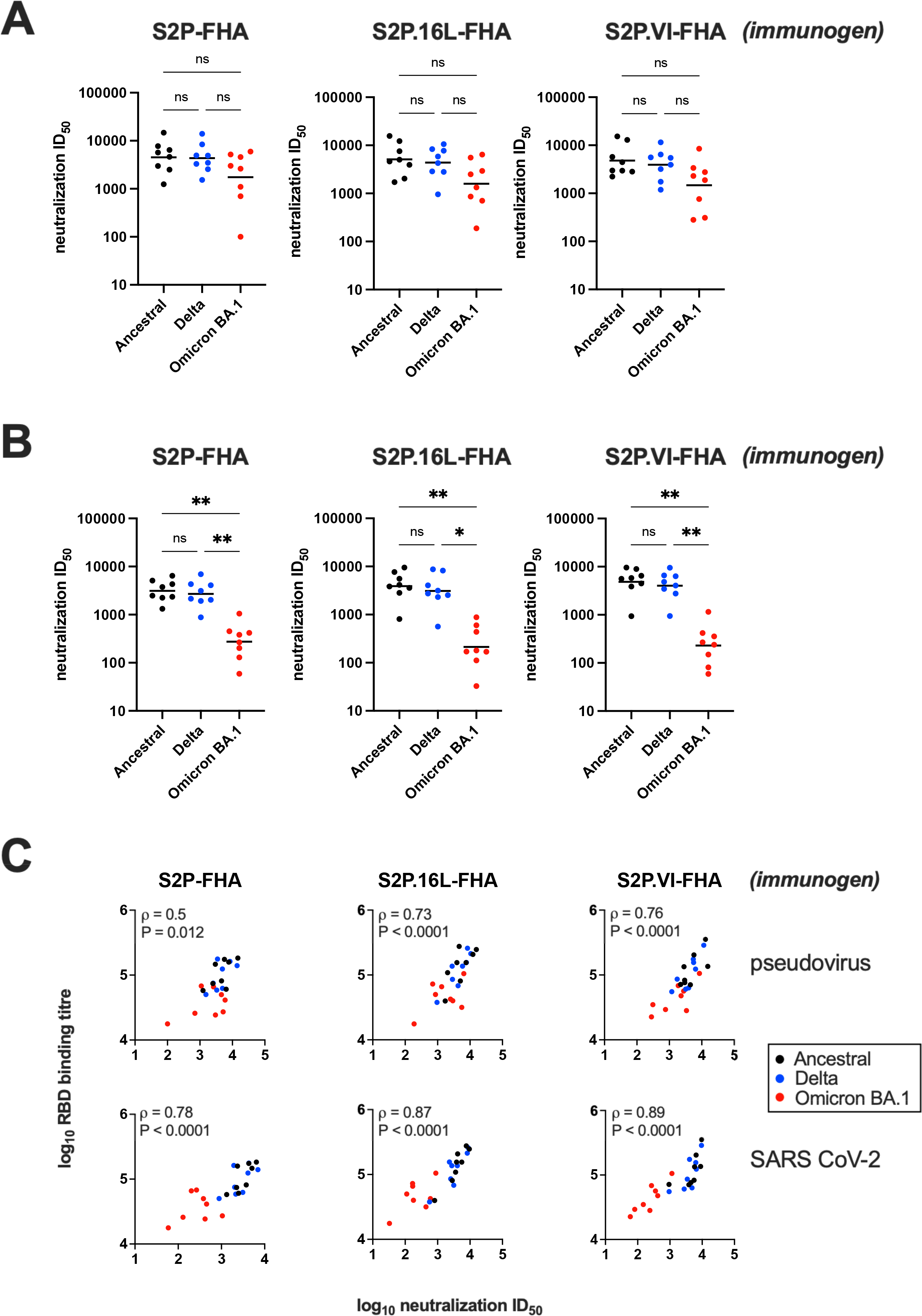
Neutralization activity of immune sera. **A)** Pseudovirus neutralization ID_50_s of week-16 vaccinal sera. The S variant genotypes used in S-HIV pseudotype assays are shown below the x axis. Neutralization ID_50_ obtained with sera from individual animals are indicated with various symbols. The bars are geometric mean ID50s. The geometric mean pseudovirus neutralization ID_50_ of the control group receiving 3 doses of 50% Addavax-PBS was <1/200. **B)** Neutralisation assays performed in high-throughput format with authentic SARS-CoV-2 variants. The SARS-CoV-2 variants are shown below the x axis. Horizontal bars are the geometric mean ID_50_s for each immunogen group. The geometric mean neutralization ID_50_ of the control group receiving 3 doses of 50% Addavax-PBS was <1/320 for Ancestral and Delta and <1/40 for Omicron BA.1. A Kruskal-Wallis test was used to determine whether the differences in ID_50_s observed between variants was significant: ns, not significant; *, P < 0.05; **, P < 0.01. **C)** Correlations between RBD binding titer (obtained in Fig. 5C) and pseudovirus neutralization ID_50_ (top) and authentic virus ID_50_ (bottom). The data points are color-coded according to variant. Spearman *ρ* and P values were determined using GraphPad Prism v9.3.0.

The neutralizing activity of sera against authentic ancestral, Delta, and Omicron BA.1 SARS-CoV-2 using HAT-24 cells in the R-20 microneutralization assay developed by Aggarwal et al. (32) was examined. Whereas ancestral clade A2.2 and Delta viruses were potently neutralized by sera from the 3 immunogen groups, Omicron BA.1 virus neutralization was reduced by ~1log_10_ (**Fig. 6B**). In **Fig. 6C**, log_10_ RBD binding titers were plotted against log_10_ neutralization ID_50_ for individual sera and color-coded according to SARS CoV-2 variant. Strong and highly significant correlations between RBD binding and authentic SARS CoV-2 neutralization (and to a lesser extent pseudoviral neutralization) were observed suggesting that RBD-directed antibodies play an important role in SARS CoV-2 neutralization. This analysis suggests that the 15 mutations in the RBD of Omicron BA.1 contribute to the decreased neutralization potency of ancestral S2P-FHA-elicited sera for Omicron BA.1 virus.

### Specificity of antibody responses

A serum-mNAb cross-competition assay was employed to gain an understanding of the antigenic sites within S that are targeted by vaccinal antibodies. Biotinylated ancestral Hu-1 S2P-FHA captured onto streptavidin coated ELISA plates was incubated with a mixture comprising a sub-saturating amount of human mNAbs or ACE2-Fc and a dilution series of sera. The assay was developed with anti-human F(ab’)2-HRP for the mNAbs or anti-human IgG-HRP for ACE2-Fc. The data **(S2 Fig)** show that S2P-FHA-, S2P.16L-FHA- and S2P.VI-FHA-elicited sera contained antibody specificities with largely equivalent abilities to block binding by ACE2-Fc and NAbs directed to the ACE2-binding site (COVOX222, SE12), a NAb directed to the NTD (COVA2-17), the stem region of S2 (CV3-25) and an undefined epitope in S (COVA1-25). The data indicate that broad specificity antibody responses were elicited by the 3 S2P-FHA antigens.

### The VI mutation confers stability to the Omicron BA.1 S2P-FHA trimer

Thirty mutations occur in the Omicron BA.1 Spike. Eight are in the NTD and 15 in the RBD, consistent with decreased neutralization of Omicron BA.1 by human vaccinee sera (32–35, 53) and low vaccine efficacy against Omicron infection (29). One approach being taken to improve the efficacy of COVID-19 vaccines against Omicron subvariants is to include Omicron-derived sequences in second-generation vaccines (36, 37). This prompted an assessment of whether the Omicron BA.1 Spike could be stabilized by mutations in the alanine cavity. Omicron BA.1 versions of S2P-FHA and S2P.VI-FHA constructs were prepared and are referred to as S2P.OmiBA1-FHA and S2P.OmiBA1.VI-FHA, respectively. The His^681^ArgAlaArgArg furin site was changed to Pro^681^GlySerAlaSer in both constructs.

The glycoproteins were expressed in Expi293F cells, extracted by divalent cation affinity chromatography and further purified by Superose 6 SEC. S2P.OmiBA1-FHA eluted as a major peak close to the position of thyroglobulin (669 kDa) (**Fig 7A**, top panel) and had an almost identical profile to that of S2P-FHA derived from the Hu-1 isolate (**see Fig 3**). The trimer was collected as a homogenous protein as indicated by analytical SEC (**Fig. 7B**, top panel). S2P.OmiBA1.VI-FHA eluted as 4 species including a prominent presumed trimer peak (dashed box, **Fig. 7A**, bottom panel). Fractions corresponding to trimer were pooled, concentrated and re-chromatographed following a freeze (−80°C)-thaw cycle in a Superose 6 column, revealing a largely homogeneous species corresponding to trimer (**Fig. 7B**). A thermofluor assay indicated that the S2P.OmiBA1-FHA trimer possessed higher thermal stability relative to its ancestral Hu-1 counterpart with a melting temperature of 61°C versus 43.6°C for the latter (**Fig. 7C**, top panel, compared with Fig. 3C top panel). The introduction of VI increased the melting temperature to 65°C for S2P.OmiBA1.VI-FHA (**Fig. 7C**, lower panel).

**Fig. 7.**
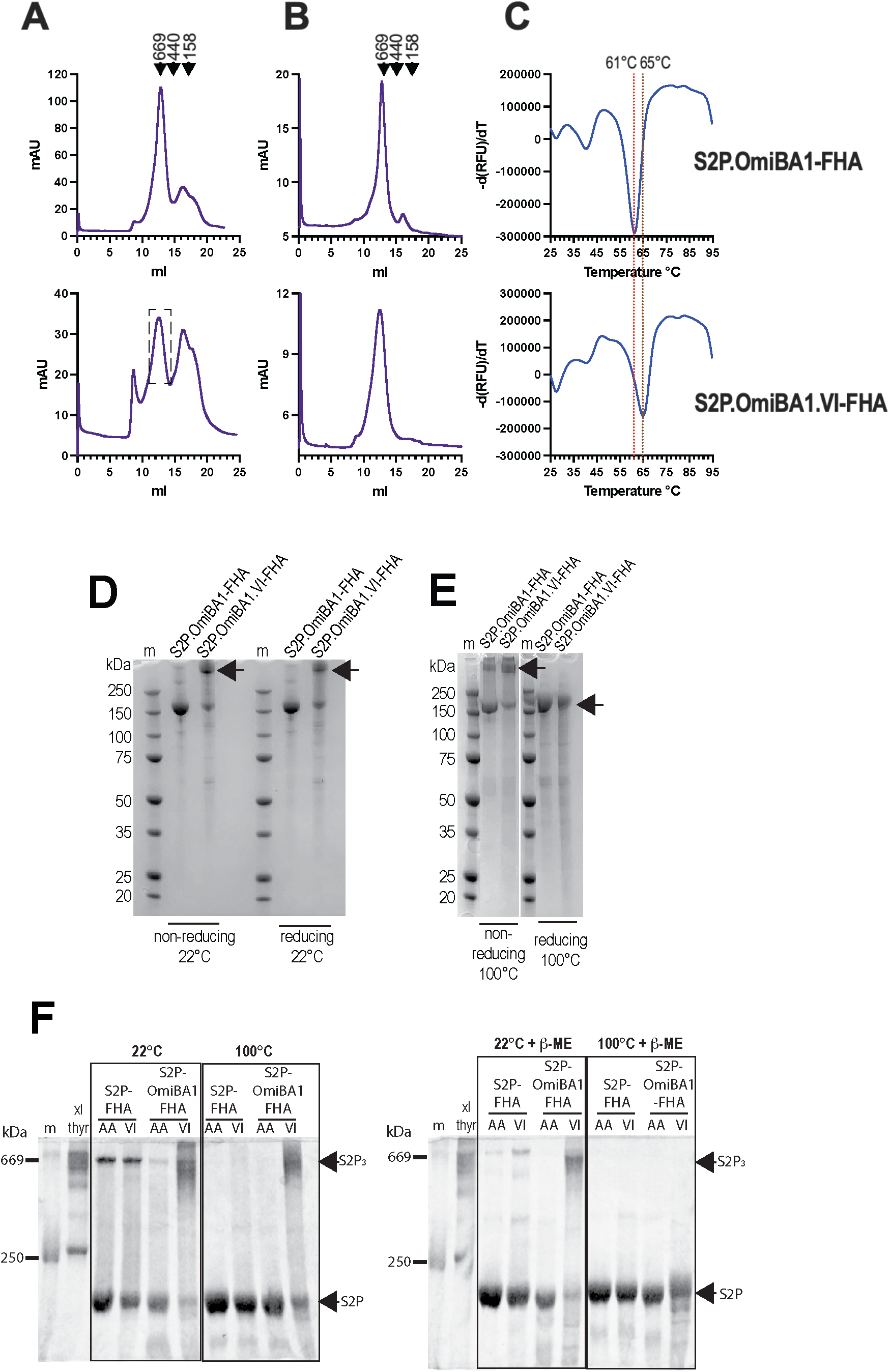
VI stabilizes S2P-FHA trimers derived from Omicron BA.1. **A**, Superose 6 SEC profiles of S2P.OmiBA1-FHA and, S2P.OmiBA1.VI-FHA eluted from TALON or HiTRAP columns, respectively. **B**, Superose 6 SEC profiles of presumed trimers obtained in **A**. **C**, Differential scanning fluorimetry of purified S2P.OmiBA1-FHA and, S2P.OmiBA1.VI-FHA trimers. **D**, SDS-PAGE/Coomassie blue staining of purified trimers under non-reducing and reducing (1% v/v betamercaptoethanol) conditions. The samples were not boiled prior to electrophoresis. m, markers. **E**, SDS-PAGE of trimers under non-reducing and reducing (1% v/v betamercaptoethanol) conditions after boiling for 5 min. The position of the S2P.omicron.VI-FHA major band under the various conditions is indicated with an arrow. m, markers. **F**, SDS-PAGE of S2P-FHA, S2P.OmiBA1-FHA, S2P.Omi.VI-FHA trimers under non-reducing (left) and reducing (1% v/v betamercaptoethanol) (right) conditions. Sample buffer containing SDS was added to the samples with and without 1% betamercaptoethanol and the samples were either left at room temperature (22°C) or boiled (100°C) for 3 min prior to electrophoresis. Thyroglobulin that had been chemically crosslinked with 1 mM bis(sulfosuccinimidyl)suberate was included to mark the expected position (669 kDa) of the S2P.OmiBA1.VI-FHA trimer (S2P_3_).

The purified S2P.OmiBA1-FHA and S2P.OmiBA1.VI-FHA trimers were further analysed by SDS-PAGE under non-reducing and reducing (1% betamercaptoethanol) conditions in the absence of sample boiling (**Fig. 7D**). The S2P.OmiBA1-FHA trimer was largely resolved to its expected monomer molecular weight of ~ 160-180 kDa in both the presence and absence of reducing agent. By contrast, the major S2P.OmiBA1.VI-FHA species (indicated by an arrow) was retained as a high molecular weight species migrating close to the top of the gel; a minor species was also observed at the monomer position. **Figure 7E** shows a repeat experiment in which the samples were boiled for 5’ prior to electrophoresis. Under non-reducing conditions, the results seen in **Figure 7D** were largely recapitulated with the high molecular weight major S2P.OmiBA1.VI-FHA species again observed. However, boiling in the presence of reducing agent resolved this species to its monomer molecular weight. The data suggest that the S2P.OmiBA1.VI-FHA trimer resists disruption by 0.8% w/v sodium dodecyl sulfate denaturant with and without 1% betamercaptoethanol at 22°C. To obtain an SDS-PAGE marker with a theoretical mol.wt of 669 kDa, which is close to that of the S2P-FHA trimer in SEC, thyroglobulin was covalently crosslinked with bis(sulfosuccinimidyl) suberate. **Figure 7F** shows that crosslinked thyroglobulin co-migrated with the SDS/betamercaptoethanol-resistant high molecular weight form of S2P.OmiBA1.VI-FHA following treatment with 0.8% SDS at 22°C or 100°C for 3 min, or 0.8% SDS + 1% betamercaptoethanol at 22°C for 3 min prior to electrophoresis. Again, S2P.OmiBA1.VI-FHA resolved as a monomer following boiling in 0.8% SDS + 1% betamercaptoethanol. S2P.OmiBA1-FHA and Hu-1 S2P-FHA proteins migrated as monomers following all treatments except for some residual trimeric S2P-FHA after treatment with 1% SDS at 22°C. The data illustrate the trimer stabilizing effect of the VI mutation.

### Antigenic properties of Ala cavity mutants

Next, biolayer interferometry was used to compare the effects of VI on the exposure of epitopes recognized by broadly neutralizing human monoclonal antibodies and ACE2-Fc in S2P-FHA trimers derived from ancestral Hu-1 and Omicron BA.1. ACE2-Fc and the human NAbs were attached to anti-human IgG Fc capture biosensors while the S2P trimers were in the analyte phase. The sensograms shown in **S3 Fig** correlate with the 1:1 bimolecular interaction model with R^2^ values being >0.95 and χ2 values being <1 for all cases (**Table 1**). A comparison of dissociation rates (k_diss_) showed that the introduction of VI to the ancestral S2P-FHA stabilized the interaction with mNAbs directed to the RBD (S2E12, S2H97, COVOX222), to the S2 stalk (CV3-25) and to an unknown epitope in S (COVA1-25) which translated to a greater-than 3log_10_ improvement in equilibrium binding constant (**Table 1**). This effect of the VI mutation was not observed in the context of S2P.OmiBA1-FHA. The data suggest that the improved thermostability of the ancestral S2P.VI-FHA trimer is associated with stabilization of its interaction with key mNAbs, however, this is not observed with its Omicron BA.1-derived counterpart.

**Table 1.** Binding kinetics of Spike trimers to ACE2-Fc and human monoclonal NAbs

| Ligand | S2P glycoprotein | R<sup>2a</sup> | χ<sup>2b</sup> | KD (M) | k<sub>on</sub> (1/Ms) | k<sub>dis</sub> (1/s) | Ligand | S2P glycoprotein | R<sup>2</sup> | χ<sup>2</sup> | KD (M) | k<sub>on</sub> (1/Ms) | k<sub>dis</sub> (1/s) |
| --- | --- | --- | --- | --- | --- | --- | --- | --- | --- | --- | --- | --- | --- |
| **S2P-FHA** |  |  |  |  |  |  | **S2P.OmiBA1-FHA** |  |  |  |  |  |  |
| **ACE2-Fc** | S2P | 0.994 | 0.056 | 1.65x10<sup>-9</sup> | 4.25x10<sup>5</sup> | 7.00x10<sup>-4</sup> | **ACE2-Fc** | S2P.OmiBA1 | 0.994 | 0.104 | 6.41x10<sup>-10</sup> | 4.20x10<sup>5</sup> | 2.69 x10<sup>-4</sup> |
|  | S2P.VI | 0.995 | 0.017 | 4.22x10<sup>-9</sup> | 1.02x10<sup>5</sup> | 4.30x10<sup>-4</sup> |  | S2P.OmiBA1.VI | 0.998 | 0.028 | 1.87x10<sup>-10</sup> | 2.74x10<sup>5</sup> | 5.10x10<sup>-5</sup> |
| **S2E12** | S2P | 0.993 | 0.316 | 2.24x10<sup>-9</sup> | 5.01x10<sup>5</sup> | 1.12x10<sup>-3</sup> | **S2E12** | S2P.OmiBA1 | 0.987 | 0.971 | 1.90x10<sup>-9</sup> | 1.29x10<sup>6</sup> | 2.44 x10<sup>-3</sup> |
|  | S2P.VI | 0.989 | 0.321 | <10<sup>-12c</sup> | 2.35x10<sup>5</sup> | <10<sup>-7</sup> |  | S2P.Omi.BA1.VI | 0.990 | 0.989 | 1.91x10<sup>-9</sup> | 8.33x10<sup>5</sup> | 1.59x10<sup>-3</sup> |
| **S2H97** | S2P | 0.981 | 1.123 | 2.03x10<sup>-10</sup> | 6.06x10<sup>5</sup> | 1.23x10<sup>-4</sup> | **S2H97** | S2P.OmiBA1 | 0.998 | 0.190 | <10<sup>-12</sup> | 2.35x10<sup>5</sup> | <10<sup>-7</sup> |
|  | S2P.VI | 0.987 | 0.390 | <10<sup>-12</sup> | 2.12x10<sup>5</sup> | <10<sup>-7</sup> |  | S2P.OmiBA1.VI | 0.999 | 0.159 | <10<sup>-12</sup> | 2.56x10<sup>5</sup> | <10<sup>-7</sup> |
| **COVOX222** | S2P | 0.996 | 0.182 | 9.29x10<sup>-10</sup> | 3.18x10<sup>5</sup> | 2.95x10<sup>-4</sup> | **COVOX222** | S2P.OmiBA1 | 0.997 | 0.215 | 4.69x10<sup>-10</sup> | 2.74x10<sup>5</sup> | 1.29x10<sup>-4</sup> |
|  | S2P.VI | 0.980 | 0.176 | <10<sup>-12</sup> | 1.05x10<sup>5</sup> | <10<sup>-7</sup> |  | S2P.OmiBA1.VI | 0.999 | 0.039 | 7.15x10<sup>-10</sup> | 1.80x10<sup>5</sup> | 1.29x10<sup>-4</sup> |
| **COVA1-25** | S2P | 0.981 | 0.015 | 1.46x10<sup>-9</sup> | 6.09x10<sup>5</sup> | 8.89x10<sup>-4</sup> | **COVA1-25** | S2P.OmiBA1 | 0.974 | 0.019 | 2.46x10<sup>-9</sup> | 3.72x10<sup>5</sup> | 9.17 x10<sup>-4</sup> |
|  | S2P.VI | 0.970 | 0.017 | <10<sup>-12</sup> | 1.06x10<sup>5</sup> | <10<sup>-7</sup> |  | S2P.OmiBA1.VI | 0.958 | 0.038 | 3.16x10<sup>-9</sup> | 4.05x10<sup>5</sup> | 1.28 x10<sup>-3</sup> |
| **CV3-25** | S2P | 0.986 | 0.359 | 2.48x10<sup>-10</sup> | 4.23x10<sup>5</sup> | 1.05x10<sup>-4</sup> | **CV3-25** | S2P.OmiBA1 | 0.999 | 0.036 | <10<sup>-12</sup> | 1.41x10<sup>5</sup> | <10<sup>-7</sup> |
|  | S2P.VI | 0.990 | 0.073 | <10<sup>-12</sup> | 1.33x10<sup>5</sup> | <10<sup>-7</sup> |  | S2P.OmiBA1.VI | 0.999 | 0.015 | 2.63x10<sup>-10</sup> | 1.32x10<sup>5</sup> | 3.5x10<sup>-5</sup> |
| **S2P.OmiBA1-1208.H8** |  |  |  |  |  |  | **S2P.OmiBA4/5-1208.H8** |  |  |  |  |  |  |
| **ACE2-Fc** | S2P.OmiBA1 | 0.995 | 0.034 | 2.15x10<sup>-9</sup> | 10<sup>6</sup> | 2.15x10<sup>-3</sup> | **ACE2-Fc** | S2P.OmiBA4/5 | 0.979 | 0.020 | 2.47x10<sup>-9</sup> | 6.25x10<sup>5</sup> | 1.54x10<sup>-3</sup> |
|  | S2P.OmiBA1.VI | 0.997 | 0.020 | 2.31x10<sup>-9</sup> | 3.16x10<sup>5</sup> | 7.31x10<sup>-4</sup> |  | S2P.OmiBA4/5.VI | 0.995 | 0.016 | 3.72x10<sup>-9</sup> | 2.29x10<sup>5</sup> | 8.52x10<sup>-4</sup> |
| **S2E12** | S2P.OmiBA1 | 0.962 | 0.500 | 2.06x10<sup>-9</sup> | 1.29x10<sup>6</sup> | 2.65x10<sup>-3</sup> | **S2H97** | S2P.OmiBA4/5 | 0.991 | 0.692 | 1.55x10<sup>-10</sup> | 6.97x10<sup>5</sup> | 1.08x10<sup>-4</sup> |
|  | S2P.Omi.BA1.VI | 0.977 | 0.356 | 2.38x10<sup>-9</sup> | 6.74x10<sup>5</sup> | 1.61x10<sup>-3</sup> |  | S2P.OmiBA4/5.VI | 0.999 | 0.030 | 5.27x10<sup>-10</sup> | 3.17x10<sup>5</sup> | 1.67x10<sup>-4</sup> |
| **S2H97** | S2P.OmiBA1 | 0.988 | 1.354 | 1.38x10<sup>-11</sup> | 7.07x10<sup>5</sup> | 9.79x10<sup>-6</sup> | **COVOX222** | S2P.OmiBA4/5 | 0.973 | 0.129 | 4.04x10<sup>-9</sup> | 5.76x10<sup>5</sup> | 2.33x10<sup>-3</sup> |
|  | S2P.OmiBA1.VI | 0.999 | 0.020 | 1.66x10<sup>-10</sup> | 1.62x10<sup>5</sup> | 2.69x10<sup>-5</sup> |  | S2P.OmiBA4/5.VI | 0.984 | 0.049 | 8.42x10<sup>-9</sup> | 4.51x10<sup>5</sup> | 3.8x10<sup>-3</sup> |
| **COVOX222** | S2P.OmiBA1 | 0.989 | 0.387 | 3.25x10<sup>-9</sup> | 6.88x10<sup>5</sup> | 2.24x10<sup>-3</sup> | **CV3-25** | S2P.OmiBA4/5 | 0.999 | 0.030 | 4.47x10<sup>-9</sup> | 3.05x10<sup>5</sup> | 1.36x10<sup>-3</sup> |
|  | S2P.OmiBA1.VI | 0.999 | 0.047 | 1.40x10<sup>-9</sup> | 2.24x10<sup>5</sup> | 3.15x10<sup>-4</sup> |  | S2P.OmiBA4/5.VI | 0.999 | 0.018 | 3.09x10<sup>-9</sup> | 1.77x10<sup>5</sup> | 5.48x10<sup>-4</sup> |
| **CV3-25** | S2P.OmiBA1 | 0.965 | 0.536 | 1.48x10<sup>-9</sup> | 4.96x10<sup>5</sup> | 7.36x10<sup>-4</sup> |  |  |  |  |  |  |  |
|  | S2P.OmiBA1.VI | 0.970 | 0.251 | <10<sup>-12</sup> | 2.54x10<sup>5</sup> | <10<sup>-7</sup> |  |  |  |  |  |  |  |

<sup>a</sup>Square of the coefficient of correlation between sensogram data and 1:1 bimolecular binding model
<sup>b</sup>Sum of the squared deviations: sensogram data versus 1:1 bimolecular binding model
<sup>c</sup>antibody-antigen complexes exhibiting negligible dissociation rate shaded in grey

### The VI mutation stabilizes the Omicron BA.1 and BA.4/5 S2P ectodomain in trimeric form in the absence of an external foldon trimerization domain

We next determined whether the VI mutation could stabilize the ancestral and Omicron BA.1 S ectodomains (residues 16-1208) in trimeric form in the absence of the C-terminal foldon motif. The last residue of the ectodomain, Q^1208^, was appended with GlySerGlySer-His_8_ to give S2P-1208.H8, S2P.VI-1208.H8 (derived from ancestral Hu-1) and S2P.OmiBA1-1208.H8 and S2P.OmiBA1.VI-1208.H8 (derived from Omicron BA.1). Superose 6 SEC revealed that the ancestral S2P-1208.H8 co-eluted with ferritin (440 kDa) suggesting that it was secreted from transfected 293F cells as a dimer whereas S2P.VI-1208.H8 was purified as a trimer, as indicated by its coelution with thyroglobulin (669 kDa) (**Fig. 8A**). The purified ancestral S2P.VI-1208.H8 trimer was reanalyzed by Superose 6 SEC following a freeze (−80°C)-thaw cycle revealing that ~ 50% of the S2P.VI-1208.H8 protein had dissociated to dimer and monomer (**Fig. 8B**). A thermofluor assay indicated that S2P.VI-1208.H8 comprised 2 species with melting temperatures of 43°C and 58°C, respectively (**Fig. 8C**). The data indicate that whereas VI enables the ancestral S2P ectodomain to form a trimer in the absence of an external trimerization tag, a large proportion of this trimer is unstable, dissociating into dimers and monomers after a freeze-thaw cycle.

**Fig. 8.**
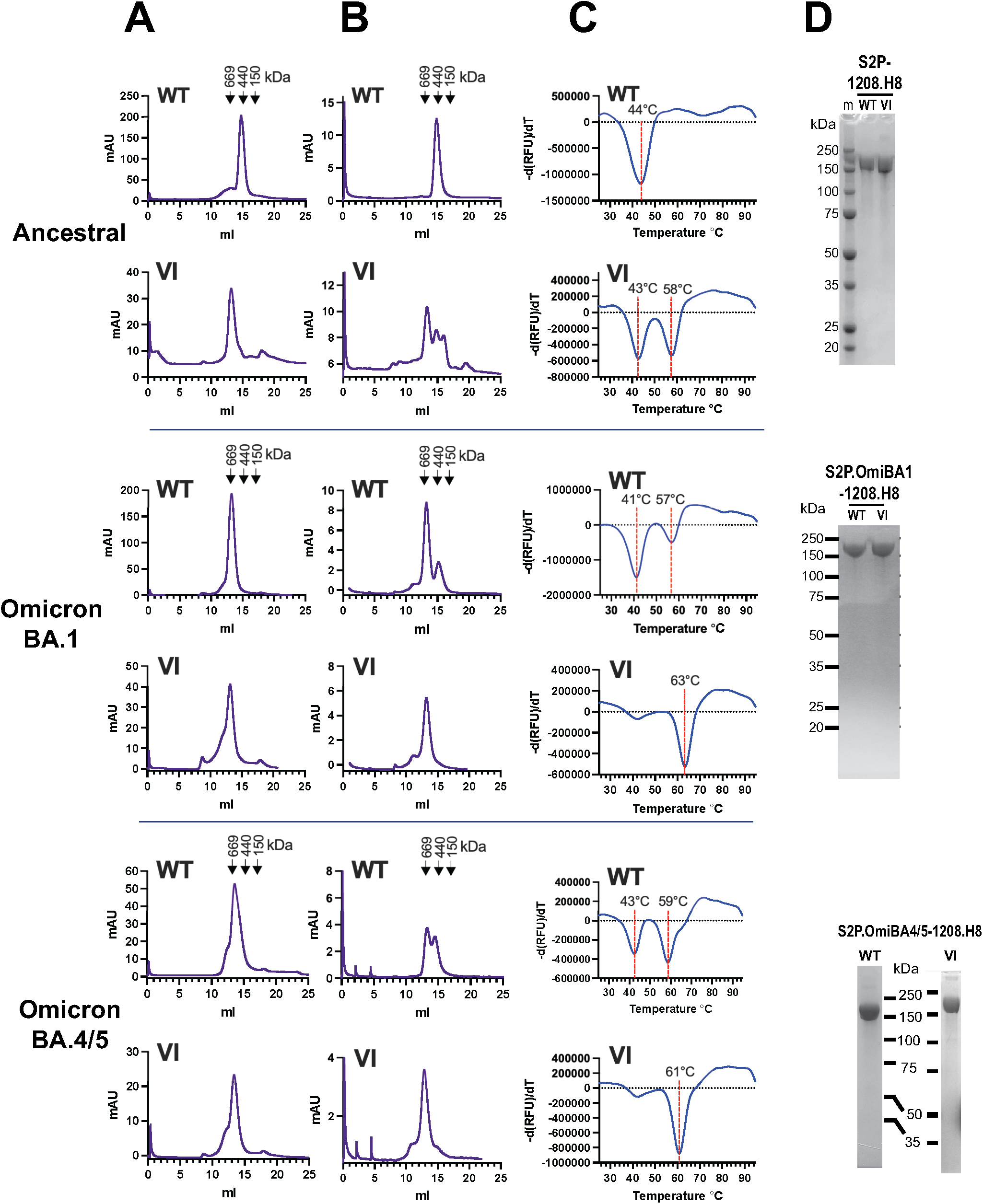
The VI mutation stabilizes the trimerization of the omicron BA.1 and BA.4/5 S ectodomain in the absence of the T4 foldon trimerization motif. **A)** Superose 6 size exclusion chromatography of S2P-1208.H8 proteins purified from 293F (ancestral) or Expi293F (Omicron BA.1 and BA.4/5) cells by hiTRAP affinity chromatography. **B**, Superose 6 size exclusion chromatography of S2P-1208.H8 glycoproteins purified in A following a freeze (−80°C)-thaw cycle. **C**, Differential scanning fluorimetry of purified S2P-1208.H8 proteins following a freeze (−80°C)-thaw cycle performed using SYPRO Orange. **D**, SDS-PAGE and Coomassie blue staining of purified proteins under reducing conditions. WT: Ala at amino positions 1016 and 1020; VI: Val and Ile at amino positions 1016 and 1020, respectively. m = markers

Both S2P.OmiBA1-1208.H8 and S2P.OmiBA1.VI-1208.H8 were obtained in trimeric form in the absence of the external foldon trimerization tag (**Fig. 8A**). Superose 6 SEC of the purified trimers following a freeze (−80°C)-thaw cycle revealed that ~ 15% of the S2P.OmiBA1-1208.H8 protein had dissociated to dimer. By contrast, the trimeric structure of S2P.Omi1.VI-1208.H8 was retained (**Fig. 8B**). A thermofluor assay indicated that the majority of purified S2P.OmiBA1-1208.H8 had a melting temperature of 41°C, whereas the melting temperature of S2P.OmiBA1.VI-1208.H8 trimers was much higher at 63°C (**Fig. 8C**). The data indicate that the S2P.OmiBA1 ectodomain is intrinsically unstable requiring the foldon domain for thermal stability. Addition of VI to the Omicron BA.1 ectodomain enables highly stable soluble trimers to be obtained thereby obviating the requirement for an exogenous trimerization domain to maintain trimeric structure.

The Omicron BA.4 and BA.5 subvariants recently evolved from the Omicron BA.2 VOC lineage and are currently the dominant variant due to an apparent transmission advantage and immune evasive properties (54). BA.4 and BA.5 S amino acid sequences are identical. In comparison to BA.1, 5/6 mutations within the NTD and 5/17 in the RBD are unique to BA.4/5; L452R and F486V within the BA.4/5 RBD are believed to be responsible for its ability to evade immunity due to vaccination and/or previous infection by Omicron BA.1 (55). The S2 subunits of the BA.4/5 spike trimer exhibit relatively tight packing, which may impose steric restrictions to NAb binding (56). We therefore prepared S2P.OmiBA.4/5-1208.H8 and S2P.OmiBA.4/5.VI-1208.H8 constructs to determine if they could also form stable trimers. The proteins were obtained largely as trimers following HiTRAP divalent affinity chromatography of transfected Expi293F culture supernatants **(Fig. 8A)**. The trimeric structure of S2P.OmiBA.4/5.VI-1208.H8 was retained following a freeze-thaw cycle, whereas a large proportion of S2P.OmiBA.4/5-1208.H8 had dissociated to putative dimer after the treatment **(Fig. 8B)**. These data were reflected in the thermofluor assay which revealed 2 species for S2P.OmiBA.4/5-1208.H8 with melting temperatures of 43°C and 59°C, whereas S2P.OmiBA.4/5.VI-1208.H8 presented as a single species with a relatively high melting temperature of 61°C **(Fig. 8C)**. Reducing SDS-PAGE and Coomassie blue staining indicated a single ~160-180kDa band for all constructs **(Fig. 8D)**. The data indicate that VI provides a method for producing soluble S glycoprotein trimers derived from Omicron subvariants in the absence of foreign, potentially immunogenic trimerization sequences.

ACE2-Fc and broadly neutralizing monoclonal antibodies were used to probe the antigenic structure of S2P.OmiBA1-1208.H8 and S2P.OmiBA.4/5-1208.H8 proteins with and without VI in biolayer interferometry. The sensograms shown in **S4 Fig** all correlate with a 1:1 bimolecular binding model with R^2^ values being >0.96 and χ2 values being <1.4 for all cases (**Table 1**). In almost all cases, the VI mutation did not substantially affect the equilibrium binding constants, association rates nor dissociation rates. The exception was the S2 stalk-directed mNAb CV3-25 which formed a highly stable interaction with S2P.OmiBA1.VI-1208.H8. S2E12 was not used for the analysis of S2P.OmiBA.4/5-1208.H8 proteins because it does not neutralize Omicron BA.4 or BA.5 S-pseudoviruses (data not shown). The data indicate that key conserved neutralization epitopes within the RBD and stalk of S2 are exposed in S2P.OmiBA1.VI-1208.H8 and S2P.OmiBA.45.VI-1208.H8 trimers.

## DISCUSSION

Class I fusion glycoproteins from a number of viral families comprise a central trimeric coiled coil that acts as a scaffold for the conformational changes required for the membrane fusion process (4, 5, 9, 38–40, 43–45, 57). In orthomyxoviruses and retroviruses, a 3-4 repeat of largely hydrophobic residues mediates knobs-into-holes interhelical contacts to help stabilize the trimer. By contrast, the coiled coil at the center of the SARS-CoV-2 prefusion S glycoprotein trimer is formed by 3 bow-shaped helices that expand away from each other from a point of contact mediated by the inward-facing Ile1013 and Leu1012. The remainder of the 3-4 repeat is largely comprised of polar residues that mediate few inter-helical contacts. This unusual structure may contribute to the relatively low thermostability of the ancestral prefusion S trimer, as evidenced by the melting temperature of 43.6°C observed here, even in the presence of the S2P mutation, shown to stabilize SARS CoV-2 S trimers in the pre-fusion conformation (5), and the C-terminal T4 foldon trimerization domain. In this study, we found that the ancestral S2P trimer could be further stabilized by the creation of an artificial hydrophobic core in the center of the coiled coil of S2 CH helices by replacing Ala1016 and Ala1020, that form part of the 3-4 repeat, with bulkier hydrophobic residues to fill the cavity associated with these Ala residues.

Mutagenesis of Ala1016 shifted the melting temperature of S2P-FHA trimers from 43.6°C to 58°C with A1016L (‘16L’) giving the highest proportion of the 58°C species. By contrast, the 43.6°C species was the predominant form with substitutions of Ala1020, including A1020L. The more pronounced stabilizing effect of the 1016 versus 1020 hydrophobic substitutions may be due to the former’s proximity to Ile1013 and substitutions at this position will enlarge the Leu1012-Ile1013 hydrophobic network that forms in the core of the coiled coil trimer. Double substitutions were all associated with the stable form, except for 1016/20VV. Interestingly, increased thermostability was generally associated with decreased soluble S2P-FHA expression suggesting that the bulk and/or geometry of the sidechain chosen to fill the cavity can impact the folding of the S2P trimer. Membrane fusion is triggered by ACE2-S1 binding and cleavage of S2 by cellular TMPRSS2, leading to viral entry and replication (4). We found that substitution of either Ala1016 or Ala1020 individually with Val or Leu. or in combination with Val and Ile, respectively, led to ~1.5-2-fold increases in membrane fusion function. The trimer stabilizing effects of filling the Ala1016/Ala1020 cavity appears linked to improved glycoprotein function. Interestingly, mutations such as A1016V, A1020V and 1020L that did not increase soluble S2P-FHA thermostability nevertheless had an enhancing effect on fusion function. Soluble S2P-FHA may therefore partially model the functional trimer with the S2 coiled coil core exhibiting subtle structural differences in the 2 forms, perhaps due to the presence of the S2P mutation or replacement of the native helical membrane anchor with foldon. Alternatively, the cavity filling mutations could impact alternative coiled coil conformers that arise during the membrane fusion cascade (9). For example, in the postfusion SARS CoV-2 S trimer, the N-terminal 2/3 of the coiled-coil is brought together in part by the trimeric packing of HR1 helices. In this conformation the 3-4 repeat residues are close enough for hydrogen bonds to form. More intimate interactions between inward facing residues of the coiled coil may cause A1016V, A1020V and A1020L to impact fusogenicity. Overall, the data indicate that creation of an artificial hydrophobic core in the center of the CH1 coiled coil of S2 improves trimer stability and membrane fusion function.

Even though the S2P.16L-FHA and S2P.VI-FHA trimers exhibited higher thermostability than unmutated S2P-FHA, the 3 antigens exhibited very similar abilities to elicit glycoprotein-binding and neutralizing antibodies against the ancestral variant (that they were derived from) as well as the Delta and Omicron BA.1 VOCs. The 3 antigens elicited very high RBD-binding titers (1/95,000-1/137,000), pseudovirus neutralizing titers (1/3,950-1/5,110) and authentic virus neutralizing titers (1/2700-1/3150) against ancestral and Delta, whereas 3-fold reductions in binding and neutralization titer were observed with RBD and pseudovirus derived from Omicron BA.1 and a more pronounced ~log_10_ reduction in neutralization titer against Omicron BA.1 authentic virus was observed (**see S1 Table**). Strong and significant correlations were observed between RBD-binding titer and authentic virus neutralizing titer suggesting that a large component of neutralizing antibody was directed to the RBD and this activity was likely compromised by the 15 mutations arising in the Omicron BA.1 RBD. The results of the serum-NAb cross-competition assay indicated that the 3 antigens elicited very similar antibody specificities that could compete with ACE2-Fc for binding to the RBM and NAbs directed to key neutralization epitopes within the RBD, NTD and S2 stem region of the ancestral S2P-FHA trimer.

The S2P-FHA trimer derived from Omicron BA.1 (S2P.OmiBA1-FHA) exhibited greater thermostability than its ancestral counterpart (melting temperature of 58°C versus 43.6°C, respectively) and introduction of VI increased the melting temperature further to 65°C. The stabilizing effects of the Ala cavity-filling VI mutation are therefore transferrable to this highly divergent variant. A comparison of the effects of VI on the antigenic structure of S2P-FHA trimers derived from ancestral, Omicron BA.1 and Omicron BA.4/5 sequences revealed that VI led to highly stable interactions between the ancestral S2P-FHA trimer to RBD-, and stem-directed mNAbs whereas such an effect was not observed with the Omicron BA.1 and BA.4/5 versions. Stabilization of the ancestral trimer by VI may cause structural changes that subtly affect access to neutralization epitopes in these regions or enable the optimization of epitope-paratope interactions such that highly stable interactions form. In theory, this may be beneficial for higher affinity interactions with B cell receptors if this construct were to be used as a vaccine immunogen. Despite these subtle changes in antigenicity, the S2P.VI-FHA trimer was as immunogenic as S2P-FHA in guinea pigs, whose B cell repertoire is unlikely to include the human mNAbs used in this study. Omicron BA.1 and BA.4/5 S trimers have been shown to have relatively compact domain organizations with increased intersubunit buried surface and relatively high stability (56, 58, 59). A more rigid Omicron BA.1 and BA.4/5 S architecture may inhibit structural changes caused by the stabilizing VI mutation in the core of the trimer such that full exposure of neutralization epitopes is maintained.

The VI mutation enabled the production of intrinsically stable Omicron BA.1 and Omicron BA.4/5 S ectodomain trimers in the absence of an external trimerization motif (S2P.OmiBA1.VI-1208.H8 and S2P.OmiBA.4/5.VI-1208.H8, respectively). The ‘foldonless’ ancestral S2P ectodomain (S2P-1208.H8) had the propensity to form dimers, requiring VI for the trimer to form. This result suggests that the sidechains of Val and Ile at positions 1016 and 1020, respectively, prefer a trimeric structure for their accommodation in the coiled coil. Despite the ancestral S2P.VI-1208.H8 ectodomain forming trimers, a large proportion dissociated to dimer and monomer following a freeze-thaw cycle indicating that the ancestral ‘foldonless’ S2P.VI-1208 trimer is unstable. By contrast, the Omicron BA.1 and Omicron BA.4/5 ‘foldonless’ ectodomains were able to form trimers in both the presence and absence of VI. However, whereas the VI mutants exhibited melting curves pointing to a single species with melting temperatures of >60°C that were stable against a freeze-thaw cycle, the non-VI versions comprised 2 species with lower melting temperatures that partially dissociated to lower-order species following freeze-thawing. The VI mutation can therefore be used to enhance the biophysical properties of candidate purified trimeric Spike glycoprotein vaccines such that foreign, highly immunogenic trimerization domains, such as T4 foldon or the HIV-1 gp41 6-helix bundle (60, 61), are no longer required. Such off target antibody responses have halted the progression of an otherwise immunogenic Spike candidate through human clinical trials (60).

The unequal global distribution of approved COVID-19 vaccines, the nondurable nature of vaccine-elicited protective NAb responses and the emergence of the SARS CoV-2 omicron lineages has led to continuing waves of SARS CoV-2 transmission. mRNA- and chimpanzee adenovirus-based COVID-19 vaccines exhibit substantial initial effectiveness against omicron BA.1, however, this wanes over time (29). NAb titers can be increased by second and third mRNA booster doses but these increases are transient (62, 63), raising the prospect of periodic boosting with circulating strain-matched vaccines to maintain protective immunity in the human population (37). Our use of hydrophobic residues to fill the Ala cavity in the core of the S trimer enabled the production of intrinsically stable omicron lineage S trimers thereby providing an avenue for developing a simple trimeric S glycoprotein vaccine that could have utility as a heterologous booster.

## MATERIALS AND METHODS

### Recombinant proteins

<u>*S2P-FHA*</u>. A synthetic gene encoding the SARS CoV-2 (ancestral Hu-1 isolate; Genbank accession number YP_009724390.1) S ectodomain, corresponding to the S2P protein described by Wrapp et al. (5), was obtained from GeneART-ThermoFisher Scientific. The gene encodes S residues 16-1208, the furin cleavage site mutation, R^682^RAR-> G^682^SAS, and a di-Pro substitution at positions 986 and 987. The C-terminus of S2P was appended with foldon (YIPEAPRDGQAYVRKDGEWVLLSTFL), octa-His and avitag (GLNDIFEAQKIEWHE) sequences, each separated by GSGS linkers to give S2P-FHA. The synthetic S2P-FHA gene was ligated downstream of a DNA sequence encoding the tissue plasminogen activator leader via *Nhe*I, within pcDNA3 (Invitrogen). Mutations were introduced into S2P-FHA expression vectors using synthetic genes encoding mutated S2P subfragments produced by GeneART-ThermoFisher Scientific. An S2P-FHA expression vector encoding the Omicron BA.1 S ectodomain was also produced by the same method (S2P.OmiBA1-FHA). <u>*RBD*</u>. Synthetic genes encoding the receptor binding domain (RBD; amino acids 332-532) of ancestral Hu-1, Delta, Omicron BA.1 and Omicron BA.4/5 isolates were obtained from GeneART-ThermoFisher Scientific and ligated to the tissue plasminogen activator leader via *Nhe*I in pcDNA3. Both proteins encode a C-terminal hexa-His tag and Avitag sequence. <u>*S2P-1208.H8*</u> expression vectors containing ancestral Hu-1, Omicron BA.1 and Omicron BA.4/5 S ectodomain sequences were derived from S2P-FHA plasmids by exchanging the foldon-His_8_-Avitag sequence with GlySerGlySer-His_8_. S2P-1208.H8 represents the S ectodomain (residues 16-1208) appended with a C-terminal His_8_ tag. <u>*hACE2-Fc*</u> is a recombinant fusion protein comprising amino acids 19-615 of the human ACE2 ectodomain linked to the Fc domain of human IgG1 via a GS linker. A synthetic gene encoding hACE2-Fc was obtained from GeneART-ThermoFisher Scientific and ligated downstream of the tissue plasminogen activator leader via *Nhe*I in pcDNA3. The DNA sequences of S and hACE2 clones were verified by fluorescent Sanger sequencing (BigDye, ABI).

### Expression and purification of recombinant proteins

S2P-FHA and S2P-1208.H8 expression vectors were transfected into 293Freestyle or Expi293F cells using 293fectin or Expifectamine, respectively, as recommended by the manufacturer (ThermoFisher Scientific). To produce biotinylated S2P-FHA and recombinant RBD proteins, the appropriate expression vectors were transfected into Expi293F-BirA cells (64) using Expifectamine. The cells were cultured for 4-5 days at 34°C after which the transfection supernatants were clarified by centrifugation and filtration through 0.45 μm nitrocellulose filters. The SARS CoV-2 glycoproteins were then purified by divalent cation affinity chromatography using TALON resin (Merck) or HiTrap IMAC FF followed by size exclusion chromatography using a Superose 6 Increase 10/300 column linked to an AKTApure instrument (Cytiva). hACE2-Fc was produced in Expi293F cells and purified from the clarified culture supernatant using Protein G-Agarose (Genscript) followed by SEC on a Superdex 200 16/600 column linked to an AKTApure instrument (Cytiva). All proteins were concentrated using Amicon centrifugal filter units. The protein solutions were filter-sterilized using 0.45 μm nitrocellulose filters and protein aliquots stored at −80°C. Protein purity was assessed by SDS-PAGE and SEC.

### Recombinant mNAbs

pCDNA3-based human IgG1 heavy and kappa and lambda light chain expression vectors (65) containing the variable regions of SARS CoV-2 directed mNAbs COVOX222 (13), S2E12 and S2H97 (18), COVA2-17 and COVA1-25 (12), and CV3-25 (23) were produced in-house using synthetic gene fragments encoding the mNAb heavy and light chain variable regions produced by GeneART-ThermoFisher Scientific. The mNAbs were produced by transfection of Expi293 cells with equal amounts of matched heavy and light chain vectors using Expifectamine according to the manufacturer’s instructions (ThermoFisher Scientific). After 5 days of culture at 37°C, the transfection supernatants were clarified by centrifugation and filtration through 0.45 μm nitrocellulose filters. The IgG was purified by affinity chromatography using Protein G-agarose (Genscript) and exchanged into PBS. The antibodies were concentrated using Amicon centrifugal filter units. IgG solutions were filter-sterilized using 0.45 μm nitrocellulose filters and aliquots stored at −80°C.

### Differential scanning fluorimetry

Differential scanning fluorimetry was used to assess protein thermostability (66). 10 μg of protein was diluted into 25μL with 5x concentration SYPRO Orange Protein Gel Stain (Sigma Aldrich) in duplicate. The samples were then heated in an Mx3005P qPCR System in 0.5°C increments from 25°C to 95°C for 1 minute per increment. 3 measurements of fluorescence were taken at the end of each increment. Excitation was at 492nm, and emission at 610nm. The melting temperature was determined to be the minimum of the negative first derivative of the melting curve.

### Cell-cell fusion assay

Mutations were introduced to the WH-Human1_EPI_402119 expression plasmid bearing codon-optimized full-length S by overlap extension PCR using Phusion DNA polymerase (ThermoFisher). The sequences of mutants were confirmed by fluorescent Sanger sequencing (BigDye Terminator v3.1, ABI). 293T effector cells were plated at 500,000 cells per well of 6-well tissue culture plates (Nunc) in Dulbecco’s modified minimal essential medium containing 10% v/v fetal bovine serum and transfected with WH-Human1_EPI_402119 plasmid (1 μg), a bacteriophage T7 RNA polymerase expression plasmid, (1 ug, pCAG-T7) (47), and a furin expression plasmid (0.25 μg, pcDNA3.1-Furin) (48). 293T-ACE2 target cells (49) in the same medium were transfected with a luciferase reporter plasmid (1 μg, pTM*luc*) (50) and a TMPRSS2 expression plasmid (0.25 μg) (51). The transfection reagent was FuGENE HD (Promega). At 48 h post transfection, effector and target cells were resuspended in fresh medium and cocultured for 3 h in round-bottomed 96-well tissue culture plates. Luciferase activity was measured using Steady-Glo luciferase reagent (Promega) in a Clariostar plate reader (BMG Labtech).

### Western blot

Transfected 293T cells were lysed for 10 min on ice in phosphate-buffered saline containing 1% Triton X-100, 0.02% sodium azide, 1 mM EDTA. Lysates were clarified by centrifugation at 10,000x*g* at 4°C prior to polyacrylamide gel electrophoresis in the presence of SDS in 10% polyacrylamide gels under reducing conditions. Proteins were transferred to nitrocellulose prior to Western blotting with anti-S1 polyclonal rabbit antibody (Sino biological) and Goat anti-rabbit IR-Dye800CW (Odyssey). The blots were scanned in a LiCORE Mol2800 (Millenium Science) and visualized using Image Studio v1.0 (LICORE).

### Biolayer interferometry

BLI-based measurements were determined using an OctetRED96 System (ForteBio, Fremont CA). Antibodies were diluted in kinetic buffer to 10 μg/ml and immobilized onto anti-human IgG Fc capture biosensors (AHC, ForteBio). Kinetics assays were carried out at 30 °C using standard kinetics acquisition rate settings (5.0 Hz, averaging by 20) at a sample plate shake speed of 1,000 rpm. The kinetic experiments included five steps: (a) baseline (180 s); (b) antibody loading (300 s); (c) second baseline (180 s); (d) association of antigen (300 s), and (e) dissociation of antigen (300 s). Fitting curves were constructed using ForteBio Data Analysis 10.0 software using a 1:1 binding model, and double reference subtraction was used for correction.

### Immunizations

Groups of 8 guinea pigs (outbred tricolor) that were matched for gender, weight, and age were immunized subcutaneously with 30 μg of S2P proteins in PBS in a 1:1 (v/v) mix with AddaVax adjuvant (InvivoGen, San Diego, CA) at weeks 0, 4 and 14. A negative control group was immunized as above with a 1:1 (v/v) mix of PBS and adjuvant. Blood was collected at 2 weeks after the 2^nd^ dose via the saphenous vein, and at 2 weeks after the 3^rd^ dose by terminal cardiac puncture and allowed to clot for serum preparation. Sera were stored at −80°C, with heat inactivation at 56 °C for 30 min prior to use in immunological assays. Animals were housed and all procedures were performed at the Preclinical, Imaging, and Research Laboratories, South Australian Health and Medical Research Institute (Gilles Plains, Australia). All animal experiments were performed in accordance with the eighth edition of the Australian Code for the Care and Use of Animals for Scientific Purposes and were approved by the SAHMRI Animal Ethics Committee, project number SAM-20-030.

### ELISA

*Streptavidin capture format*. Streptavidin capture format. Nunc Maxisorp 96 well plates were coated with streptavidin (5 μg/ml, 50 mM carbonate buffer) overnight at 4°C, washed with PBS and blocked with BSA (10 mg/ml, PBS) at room temperature for 1 h. The plates were again washed and then incubated with biotinylated RBD (5 μg/ml, PBS) for overnight at 4°C. After further washing, the plates were incubated with serially diluted serum samples for 2 h at room temperature and antibody binding detected using horseradish peroxidase–labelled rabbit anti-guinea pig antibody (Dako, Glostrup, Denmark) and 5,5’-dithiobis-(2-nitrobenzoic acid). *Direct binding format*. Nunc Maxisorp 96 well plates were coated with S2P-FHA protein solutions (2 μg/ml, PBS) at 4°C overnight. The plates were washed with PBS and blocked with BSA (10 μg/ml, PBS) at room temperature for 1 h. The plates were again washed and then incubated with serially diluted serum samples for 2 h at room temperature. Antibody binding was detected using horseradish peroxidase– labelled rabbit anti-guinea pig antibody (Dako, Glostrup, Denmark) and 5,5’-dithiobis-(2-nitrobenzoic acid). Color reactions were measured with a Clariostar plate reader (BMG Lab Technologies). Optical density was plotted against the reciprocal dilution of plasma in Prism v9.3.0 and curves fitted using the Sigmoidal, 4PL, X is concentration model. The binding titer was defined as the reciprocal dilution of serum giving an optical density ten-times that of background, as defined by binding to BSA.

### Pseudotyped virus production

S-pseudotyped HIV luciferase reporter viruses were prepared according to the method of Jackson et al. (52). Plasmids for the production of S-HIV pseudoparticles were a kind gift of Professor Doria-Rose, NIH Vaccine Research Center, and include the WH-Human1_EPI_402119 expression plasmid bearing codon-optimized full-length S (Genbank # MN908947.3), the packaging plasmid pCMVΔR8.2 and luciferase reporter plasmid pHR’ CMV Luc (67), and a TMPRSS2 plasmid (51). The 4 plasmids were co-transfected into HEK293T cells and after 18 h of incubation, the medium was replaced with fresh Dulbecco’s modification of minimal essential medium containing 10% fetal bovine serum (DMF10) and cultured for a further 2 days. Clarified supernatants containing retroviral pseudotyped viruses were filtered through 0.45μm membrane filters. WH-Human1_EPI_402119 vectors encoding synthetic Delta and Omicron BA.1 S genes produced by GeneArt were also produced.

### S-HIV pseudovirus neutralizing assay

Neutralization assays were conducted according to the method of Jackson et al. (52). Heat inactivated sera (56°C for 30 minutes) were serially diluted in DMF10 and each dilution mixed with an equal volume of S-pseudotyped HIV luciferase reporter viruses and incubated for 1h at 37°C in triplicate. Virus-serum mixtures were added to 293T-ACE2 cell (49) monolayers attached to poly-L-lysine (0.01% w/v) coated 96 well plates the day prior at 10,000 cells/well and incubated for 2h at 37°C before addition of an equal volume of DMF10. After 3 days, tissue culture fluid was removed, monolayers were washed once with PBS and lysed with cell culture lysis reagent (Promega) and luciferase measured using luciferase substrate (Promega) in a CLARIOstar plate reader (BMG LabTechnologies). The mean percentage entry was calculated as (RLU plasma+virus)/(RLU medium+virus)*100. The percentage entry was plotted against the reciprocal dilution of plasma in Prism v9.3.0 and curves fitted using the Sigmoidal, 4PL, X is concentration model. The reciprocal dilution of plasma required to prevent 50% virus entry was calculated from the non-linear regression line (ID_50_). The lowest amount of neutralizing antibody detectable is a titer of 200. All samples that did not reach 50% neutralization were assigned an arbitrary value of 100.

### Authentic virus neutralization assay

The neutralizing activity of sera against authentic ancestral hCoV-19/Australia/NSW2715/2020 (Spike sequence identical to Hu-1), Delta, and Omicron BA.1 SARS-CoV-2 was determined with the rapid high-content SARS-CoV-2 microneutralization assay described by Aggarwal et al. (32). Briefly, Hoechst-33342-stained HAT-24 cells were seeded in 384-well plates (Corning, CLS3985). Serially diluted heat-inactivated vaccinal sera were coincubated with an equal volume of SARS-CoV-2 virus solution at twice the median lethal dose for 1 h at 37 °C. 40 μl of serum-virus mixtures were added to an equal volume of pre-plated cells, incubated for 20 h and then directly imaged on an InCell Analyzer HS2500 high-content fluorescence microscopy system (Cytiva). Cellular nuclei counts were obtained with IN Carta automated image analysis software (Cytiva), and the percentage of virus neutralization was calculated as described in (32). The neutralization ID_50_ was the last consecutive dilution reaching ≥50% neutralization.

### Serum-mNAb cross-competition ELISA

Biotinylated S2P-FHA proteins were produced in Expi293F-BirA cells (Expi293F cells that stably express BirA) and purified as described above for S2P-FHA. For competition ELISAs, Nunc Maxisorp 96-well plates were coated with streptavidin (Sigma) (5 μg/ml in 50 mM carbonate buffer) at 4°C, overnight after which they were blocked with BSA (10 mg/ml in PBS) at room temperature for 1h. After 2 washes, the plates were incubated with biotinylated S2P-FHA trimers (2 μg/ml in 5 mg/ml BSA/PBS containing 0.05% Tween 20) at room temperature for 1 h. Serially diluted vaccinal sera were mixed with sub-saturating amounts of hACE2-Fc and anti-S mNAbs and incubated with the streptavidin-biotinylated S2P-FHA coated plates for a further 2 h at room temperature. mNAb binding was detected using horseradish peroxidase–labelled goat anti-human IgG F(ab’)2 (Thermofisher-Scientific), or horseradish peroxidase–labelled anti-human IgA, IgG, IgM (Dako, Glostrup, Denmark) for hACE2-Fc. The substrate was 5,5’-dithiobis-(2-nitrobenzoic acid). Color reactions were measured with a Multiskan Ascent plate reader (Thermo Electron, Waltham, MA). Antibody binding to different antigens was compared by fitting curves with nonlinear regression using Prism version 9 software, and ID50s obtained by interpolation.

### Statistical methods

Data were statistically compared using the non-parametric Kruskal-Wallis test with Dunn’s multiple comparisons in Prism 9.3.0. Correlations between RBD titer and neutralization ID50 were tested using the nonparametric Spearman test. A P value of < 0.05 was considered significant.

## ACKNOWLEDGEMENTS

293-ACE2 cells were obtained from Professor Jesse Bloom, Fred Hutch Cancer Center, Seattle, WA. The WH-Human1_EPI_402119, pCMVΔR8.2, pHR’ CMV Luc, and TMPRSS2 expression plasmids were obtained from Professor Nicole Doria-Rose, Vaccine Research Center, NIH, MD, USA. Expi293F-BirA cells were obtained from Dr. Bruce Wines, Burnet Institute, VIC, Australia.

---

**S1 Fig. Characterization of purified S2P-FHA. A**, Superose 6 SEC of purified S2P-FHA. Standards: thyroglobulin, 669 kDa, aldolase, 158 kDa. **B**, SDS-PAGE under reducing conditions and Coomassie blue staining of purified S2P-FHA. **C**, Binding of ACE2-Fc and human mNAbs to avidin-captured biotinylated S2P-FHA in ELISA. **D**, Differential scanning fluorimetry of purified S2P-FHA using SYPRO Orange. The rate of change of fluorescence over time [– d(RFU)/dt] as a function of temperature is shown.

**S2 Fig. Specificity of elicited antibodies assessed by competition ELISA**. Competition ID_50_s of vaccinal sera versus ACE2-Fc or mNAbs for binding to streptavidin-captured ancestral Hu-1 biotin-S2P-FHA. Serially diluted vaccinal sera were mixed with constant amounts of ACE2-Fc and human monoclonal anti-S IgGs prior to incubation with streptavidin captured biotin-S2P-FHA. The immunogen groups are indicated below the graphs. A Kruskal-Wallis test was used to determine that the differences in ID_50_s observed between groups was not significant (ns).

**S3 Fig. BLI measurements of ACE2-Fc and mNAbs binding to Ala cavity mutants**. Binding of S2P-FHA and S2P.VI-FHA analytes derived from ancestral Hu-1 (left) and Omicron BA.1 (right) to S ligands immobilized on anti-human IgG capture biosensors. Association was for 300 sec followed by dissociation for 300 sec.

**S4 Fig**. BLI measurements of ACE2-Fc and human monoclonal anti-S IgGs immobilized on anti-human IgG capture biosensors binding to S2P.OmiBA1-1208.H8 (WT) and S2P.OmiBA1.VI-1208.H8 (VI) in the analyte phase. Association was for 300 sec followed by dissociation for 300 sec.

## REFERENCES

1. Duan L, Zheng Q, Zhang H, Niu Y, Lou Y, Wang H. The SARS-CoV-2 Spike Glycoprotein Biosynthesis, Structure, Function, and Antigenicity: Implications for the Design of Spike-Based Vaccine Immunogens. Front Immunol. 2020;11:576622.

2. Finkelstein MT, Mermelstein AG, Parker Miller E, Seth PC, Stancofski ED, Fera D. Structural Analysis of Neutralizing Epitopes of the SARS-CoV-2 Spike to Guide Therapy and Vaccine Design Strategies. Viruses. 2021;13(1).

3. Hoffmann M, Kleine-Weber H, Pohlmann S. A Multibasic Cleavage Site in the Spike Protein of SARS-CoV-2 Is Essential for Infection of Human Lung Cells. Mol Cell. 2020;78(4):779–84 e5.

4. Walls AC, Park YJ, Tortorici MA, Wall A, McGuire AT, Veesler D. Structure, Function, and Antigenicity of the SARS-CoV-2 Spike Glycoprotein. Cell. 2020;183(6):1735.

5. Wrapp D, Wang N, Corbett KS, Goldsmith JA, Hsieh CL, Abiona O, et al. Cryo-EM structure of the 2019-nCoV spike in the prefusion conformation. Science. 2020;367(6483):1260–3.

6. Fu Q, Chou JJ. A Trimeric Hydrophobic Zipper Mediates the Intramembrane Assembly of SARS-CoV-2 Spike. J Am Chem Soc. 2021;143(23):8543–6.

7. Ke Z, Oton J, Qu K, Cortese M, Zila V, McKeane L, et al. Structures and distributions of SARS-CoV-2 spike proteins on intact virions. Nature. 2020;588(7838):498–502.

8. Jackson CB, Farzan M, Chen B, Choe H. Mechanisms of SARS-CoV-2 entry into cells. Nat Rev Mol Cell Biol. 2022;23(1):3–20.

9. Cai Y, Zhang J, Xiao T, Peng H, Sterling SM, Walsh RM, Jr., et al. Distinct conformational states of SARS-CoV-2 spike protein. Science. 2020;369(6511):1586–92.

10. Piccoli L, Park YJ, Tortorici MA, Czudnochowski N, Walls AC, Beltramello M, et al. Mapping Neutralizing and Immunodominant Sites on the SARS-CoV-2 Spike Receptor-Binding Domain by Structure-Guided High-Resolution Serology. Cell. 2020;183(4):1024–42 e21.

11. Barnes CO, West AP, Jr., Huey-Tubman KE, Hoffmann MAG, Sharaf NG, Hoffman PR, et al. Structures of Human Antibodies Bound to SARS-CoV-2 Spike Reveal Common Epitopes and Recurrent Features of Antibodies. Cell. 2020;182(4):828–42 e16.

12. Brouwer PJM, Caniels TG, van der Straten K, Snitselaar JL, Aldon Y, Bangaru S, et al. Potent neutralizing antibodies from COVID-19 patients define multiple targets of vulnerability. Science. 2020;369(6504):643–50.

13. Dejnirattisai W, Zhou D, Supasa P, Liu C, Mentzer AJ, Ginn HM, et al. Antibody evasion by the P.1 strain of SARS-CoV-2. Cell. 2021;184(11):2939–54 e9.

14. Gruell H, Vanshylla K, Weber T, Barnes CO, Kreer C, Klein F. Antibody-mediated neutralization of SARS-CoV-2. Immunity. 2022;55(6):925–44.

15. Liu L, Wang P, Nair MS, Yu J, Rapp M, Wang Q, et al. Potent neutralizing antibodies against multiple epitopes on SARS-CoV-2 spike. Nature. 2020;584(7821):450–6.

16. Qi H, Liu B, Wang X, Zhang L. The humoral response and antibodies against SARS-CoV-2 infection. Nat Immunol. 2022;23(7):1008–20.

17. Shi R, Shan C, Duan X, Chen Z, Liu P, Song J, et al. A human neutralizing antibody targets the receptor-binding site of SARS-CoV-2. Nature. 2020;584(7819):120–4.

18. Starr TN, Czudnochowski N, Liu Z, Zatta F, Park YJ, Addetia A, et al. SARS-CoV-2 RBD antibodies that maximize breadth and resistance to escape. Nature. 2021;597(7874):97–102.

19. Andreano E, Piccini G, Licastro D, Casalino L, Johnson NV, Paciello I, et al. SARS-CoV-2 escape from a highly neutralizing COVID-19 convalescent plasma. Proc Natl Acad Sci U S A. 2021;118(36).

20. Cerutti G, Guo Y, Zhou T, Gorman J, Lee M, Rapp M, et al. Potent SARS-CoV-2 neutralizing antibodies directed against spike N-terminal domain target a single supersite. Cell Host Microbe. 2021;29(5):819–33 e7.

21. McCallum M, De Marco A, Lempp FA, Tortorici MA, Pinto D, Walls AC, et al. N-terminal domain antigenic mapping reveals a site of vulnerability for SARS-CoV-2. Cell. 2021;184(9):2332–47 e16.

22. McCarthy KR, Rennick LJ, Nambulli S, Robinson-McCarthy LR, Bain WG, Haidar G, et al. Recurrent deletions in the SARS-CoV-2 spike glycoprotein drive antibody escape. Science. 2021;371(6534):1139–42.

23. Jennewein MF, MacCamy AJ, Akins NR, Feng J, Homad LJ, Hurlburt NK, et al. Isolation and characterization of cross-neutralizing coronavirus antibodies from COVID-19+ subjects. Cell Rep. 2021;36(2):109353.

24. Li W, Chen Y, Prevost J, Ullah I, Lu M, Gong SY, et al. Structural basis and mode of action for two broadly neutralizing antibodies against SARS-CoV-2 emerging variants of concern. Cell Rep. 2022;38(2):110210.

25. Chen KK, Tsung-Ning Huang D, Huang LM. SARS-CoV-2 variants - Evolution, spike protein, and vaccines. Biomed J. 2022;45(4):573–9.

26. Ghimire D, Han Y, Lu M. Structural Plasticity and Immune Evasion of SARS-CoV-2 Spike Variants. Viruses. 2022;14(6).

27. Jacobs JL, Haidar G, Mellors JW. COVID-19: Challenges of Viral Variants. Annu Rev Med. 2022.

28. Li J, Jia H, Tian M, Wu N, Yang X, Qi J, et al. SARS-CoV-2 and Emerging Variants: Unmasking Structure, Function, Infection, and Immune Escape Mechanisms. Front Cell Infect Microbiol. 2022;12:869832.

29. Andrews N, Stowe J, Kirsebom F, Toffa S, Rickeard T, Gallagher E, et al. Covid-19 Vaccine Effectiveness against the Omicron (B.1.1.529) Variant. N Engl J Med. 2022;386(16):1532–46.

30. Lopez Bernal J, Andrews N, Gower C, Gallagher E, Simmons R, Thelwall S, et al. Effectiveness of Covid-19 Vaccines against the B.1.617.2 (Delta) Variant. N Engl J Med. 2021;385(7):585–94.

31. Supasa P, Zhou D, Dejnirattisai W, Liu C, Mentzer AJ, Ginn HM, et al. Reduced neutralization of SARS-CoV-2 B.1.1.7 variant by convalescent and vaccine sera. Cell. 2021;184(8):2201–11 e7.

32. Aggarwal A, Stella AO, Walker G, Akerman A, Esneau C, Milogiannakis V, et al. Platform for isolation and characterization of SARS-CoV-2 variants enables rapid characterization of Omicron in Australia. Nat Microbiol. 2022;7(6):896–908.

33. Dejnirattisai W, Huo J, Zhou D, Zahradnik J, Supasa P, Liu C, et al. SARS-CoV-2 Omicron-B.1.1.529 leads to widespread escape from neutralizing antibody responses. Cell. 2022;185(3):467–84 e15.

34. Nutalai R, Zhou D, Tuekprakhon A, Ginn HM, Supasa P, Liu C, et al. Potent cross-reactive antibodies following Omicron breakthrough in vaccinees. Cell. 2022;185(12):2116–31 e18.

35. Hachmann NP, Miller J, Collier AY, Ventura JD, Yu J, Rowe M, et al. Neutralization Escape by SARS-CoV-2 Omicron Subvariants BA.2.12.1, BA.4, and BA.5. N Engl J Med. 2022;387(1):86–8.

36. Branche AR, Rouphael NG, Diemert DJ, Falsey AR, Losada C, Baden LR, et al. SARS-CoV-2 Variant Vaccine Boosters Trial: Preliminary Analyses. medRxiv. [Preprint] 2022. Available from: https://www.medrxiv.org/content/10.1101/2022.07.12.22277336v1 doi: 10.1101/2022.07.12.22277336

37. Chalkias S, Harper C, Vrbicky K, Walsh SR, Essink B, Brosz A, et al. A Bivalent Omicron-Containing Booster Vaccine against Covid-19. N Engl J Med. 2022;387(14):1279–91.

38. Bullough PA, Hughson FM, Skehel JJ, Wiley DC. Structure of influenza haemagglutinin at the pH of membrane fusion. Nature. 1994;371(6492):37–43.

39. Chan DC, Fass D, Berger JM, Kim PS. Core structure of gp41 from the HIV envelope glycoprotein. Cell. 1997;89(2):263–73.

40. Julien JP, Cupo A, Sok D, Stanfield RL, Lyumkis D, Deller MC, et al. Crystal structure of a soluble cleaved HIV-1 envelope trimer. Science. 2013;342(6165):1477–83.

41. McLellan JS, Yang Y, Graham BS, Kwong PD. Structure of respiratory syncytial virus fusion glycoprotein in the postfusion conformation reveals preservation of neutralizing epitopes. J Virol. 2011;85(15):7788–96.

42. McLellan JS, Chen M, Joyce MG, Sastry M, Stewart-Jones GB, Yang Y, et al. Structure-based design of a fusion glycoprotein vaccine for respiratory syncytial virus. Science. 2013;342(6158):592–8.

43. Walls AC, Tortorici MA, Snijder J, Xiong X, Bosch BJ, Rey FA, et al. Tectonic conformational changes of a coronavirus spike glycoprotein promote membrane fusion. Proc Natl Acad Sci U S A. 2017;114(42):11157–62.

44. Weissenhorn W, Dessen A, Harrison SC, Skehel JJ, Wiley DC. Atomic structure of the ectodomain from HIV-1 gp41. Nature. 1997;387(6631):426–30.

45. Wilson IA, Skehel JJ, Wiley DC. Structure of the haemagglutinin membrane glycoprotein of influenza virus at 3 A resolution. Nature. 1981;289(5796):366–73.

46. Baase WA, Liu L, Tronrud DE, Matthews BW. Lessons from the lysozyme of phage T4. Protein Sci. 2010;19(4):631–41.

47. Takikawa S, Ishii K, Aizaki H, Suzuki T, Asakura H, Matsuura Y, et al. Cell fusion activity of hepatitis C virus envelope proteins. J Virol. 2000;74(11):5066–74.

48. Binley JM, Sanders RW, Clas B, Schuelke N, Master A, Guo Y, et al. A recombinant human immunodeficiency virus type 1 envelope glycoprotein complex stabilized by an intermolecular disulfide bond between the gp120 and gp41 subunits is an antigenic mimic of the trimeric virion-associated structure. J Virol. 2000;74(2):627–43.

49. Crawford KHD, Eguia R, Dingens AS, Loes AN, Malone KD, Wolf CR, et al. Protocol and Reagents for Pseudotyping Lentiviral Particles with SARS-CoV-2 Spike Protein for Neutralization Assays. Viruses. 2020;12(5).

50. Maerz AL, Center RJ, Kemp BE, Kobe B, Poumbourios P. Functional implications of the human T-lymphotropic virus type 1 transmembrane glycoprotein helical hairpin structure [In Process Citation]. J Virol. 2000;74(14):6614–21.

51. Bottcher E, Matrosovich T, Beyerle M, Klenk HD, Garten W, Matrosovich M. Proteolytic activation of influenza viruses by serine proteases TMPRSS2 and HAT from human airway epithelium. J Virol. 2006;80(19):9896–8.

52. Jackson LA, Anderson EJ, Rouphael NG, Roberts PC, Makhene M, Coler RN, et al. An mRNA Vaccine against SARS-CoV-2 - Preliminary Report. N Engl J Med. 2020;383(20):1920–31.

53. Tada T, Zhou H, Dcosta BM, Samanovic MI, Chivukula V, Herati RS, et al. Increased resistance of SARS-CoV-2 Omicron variant to neutralization by vaccine-elicited and therapeutic antibodies. EBioMedicine. 2022;78:103944.

54. Aggarwal A, Akerman A, Milogiannakis V, Silva MR, Walker G, Stella AO, et al. SARS-CoV-2 Omicron BA.5: Evolving tropism and evasion of potent humoral responses and resistance to clinical immunotherapeutics relative to viral variants of concern. EBioMedicine. 2022;84:104270.

55. Tuekprakhon A, Nutalai R, Dijokaite-Guraliuc A, Zhou D, Ginn HM, Selvaraj M, et al. Antibody escape of SARS-CoV-2 Omicron BA.4 and BA.5 from vaccine and BA.1 serum. Cell. 2022;185(14):2422–33 e13.

56. Cao Y, Yisimayi A, Jian F, Song W, Xiao T, Wang L, et al. BA.2.12.1, BA.4 and BA.5 escape antibodies elicited by Omicron infection. Nature. 2022;608(7923):593–602.

57. Benton DJ, Gamblin SJ, Rosenthal PB, Skehel JJ. Structural transitions in influenza haemagglutinin at membrane fusion pH. Nature. 2020;583(7814):150–3.

58. Cerutti G, Guo Y, Liu L, Liu L, Zhang Z, Luo Y, et al. Cryo-EM structure of the SARS-CoV-2 Omicron spike. Cell Rep. 2022;38(9):110428.

59. Cui Z, Liu P, Wang N, Wang L, Fan K, Zhu Q, et al. Structural and functional characterizations of infectivity and immune evasion of SARS-CoV-2 Omicron. Cell. 2022;185(5):860–71 e13.

60. Chappell KJ, Mordant FL, Li Z, Wijesundara DK, Ellenberg P, Lackenby JA, et al. Safety and immunogenicity of an MF59-adjuvanted spike glycoprotein-clamp vaccine for SARS-CoV-2: a randomised, double-blind, placebo-controlled, phase 1 trial. Lancet Infect Dis. 2021;21(10):1383–94.

61. Sliepen K, van Montfort T, Melchers M, Isik G, Sanders RW. Immunosilencing a highly immunogenic protein trimerization domain. J Biol Chem. 2015;290(12):7436–42.

62. Bar-On YM, Goldberg Y, Mandel M, Bodenheimer O, Amir O, Freedman L, et al. Protection by a Fourth Dose of BNT162b2 against Omicron in Israel. N Engl J Med. 2022;386(18):1712–20.

63. Pajon R, Doria-Rose NA, Shen X, Schmidt SD, O’Dell S, McDanal C, et al. SARS-CoV-2 Omicron Variant Neutralization after mRNA-1273 Booster Vaccination. N Engl J Med. 2022;386(11):1088–91.

64. Wines BD, Vanderven HA, Esparon SE, Kristensen AB, Kent SJ, Hogarth PM. Dimeric FcgammaR Ectodomains as Probes of the Fc Receptor Function of Anti-Influenza Virus IgG. J Immunol. 2016;197(4):1507–16.

65. Center RJ, Boo I, Phu L, McGregor J, Poumbourios P, Drummer HE. Enhancing the antigenicity and immunogenicity of monomeric forms of hepatitis C virus E2 for use as a preventive vaccine. J Biol Chem. 2020;295(21):7179–92.

66. Niesen FH, Berglund H, Vedadi M. The use of differential scanning fluorimetry to detect ligand interactions that promote protein stability. Nat Protoc. 2007;2(9):2212–21.

67. Naldini L, Blomer U, Gage FH, Trono D, Verma IM. Efficient transfer, integration, and sustained long-term expression of the transgene in adult rat brains injected with a lentiviral vector. Proc Natl Acad Sci U S A. 1996;93(21):11382–8.

